# Planar, Spiral, and Concentric Traveling Waves Distinguish Cognitive States in Human Memory

**DOI:** 10.1101/2024.01.26.577456

**Authors:** Anup Das, Erfan Zabeh, Bard Ermentrout, Joshua Jacobs

## Abstract

A fundamental challenge in neuroscience is explaining how widespread brain regions flexibly interact to support behaviors. We hypothesize that traveling waves of oscillations are a key mechanism of neural coordination, such that they propagate across the cortex in distinctive patterns that control how different regions interact. To test this hypothesis, we used direct brain recordings from humans performing multiple memory experiments and an analytical framework that flexibly measures the propagation patterns of traveling waves. We found that traveling waves propagated along the cortex in not only plane waves, but also spirals, sources and sinks, and more complex patterns. The propagation patterns of traveling waves correlated with novel aspects of behavior, with specific wave shapes reflecting particular cognitive processes and even individual remembered items. Our findings suggest that large-scale cortical patterns of traveling waves reveal the spatial organization of cognitive processes in the brain and may be relevant for neural decoding.

## Introduction

Neurons in the human cortex have rich dendritic arbors that integrate diverse inputs and axons that project outputs to multiple, distributed areas (Swanson, 2003). However, recent developments in brain imaging have suggested a dynamic interplay of distributed brain regions that underlie complex human behaviors (Sporns et al., 2004). How do individual neurons or regions reorganize their activity so that they are selectively and dynamically linked to certain other areas even as the structure of these neurons does not change on the timescale of behavior? We hypothesize that propagating patterns of brain oscillations, or “traveling waves,” underlie this selective reorganization. Traveling waves propagate across the cortex in specific directions during behavior, so that certain spatial arrangements of neurons are active at similar times, thus flexibly and efficiently communicating between the given brain regions that are relevant for a task (Mohan et al., 2024). Traveling waves are present at various frequencies (Ermentrout & Kleinfeld, 2001), including the theta/alpha range in humans (Zhang et al., 2018). Because propagating oscillations correlate with underlying neuronal activity (Jacobs et al., 2007), the spatiotemporal organization of traveling waves at each moment may indicate which brain areas are active and how they correspond to particular computational processes and memory representations in the cortex (Pinotsis et al., 2023; Pinotsis & Miller, 2023). The propagation of traveling waves may therefore show how the brain’s spatial organization and connectivity flexibly adapts to task demands and behaviors.

Traveling waves are known to play a critical role for behaviors such as visual processing and spatial navigation in rodents and non-human primates (Agarwal et al., 2014; Davis et al., 2021; Lubenov & Siapas, 2009; Muller et al., 2014; Zanos et al., 2015), and recently in human cognition (Alamia & VanRullen, 2019; Kleen et al., 2021; Mahjoory et al., 2020; Stolk et al., 2019; Zhang et al., 2018). Traveling waves also appear in spirals during sleep in humans (Muller et al., 2016). However, our current understanding of traveling waves in cognition is largely limited to planar waves, where a consistent direction of wave propagation is maintained across a large region of cortex. It is possible that other types of traveling waves with more complex spatial patterns are also relevant functionally (Bhattacharya, Brincat, et al., 2022; Denker et al., 2018; Freeman, 2009; Muller et al., 2016), so this focus on plane waves in humans may have limited our understanding of the role of traveling waves and oscillations in modulating cortical representations to support behavior.

We recently found that the direction of traveling wave propagation during behavior correlated with distinct memory processes (Mohan et al., 2024). Extending this idea, we tested here the broader idea that the spatial patterning of wave propagation changed shape based on a person’s current behavior. We designed a flexible analytical framework for measuring general patterns of traveling wave propagation and applied this procedure to direct brain recordings from neurosurgical patients performing multiple memory experiments. Our results showed an array of complex traveling wave propagation patterns including spirals, concentric waves (sources and sinks) (also known as target waves (Hagan, 1981; Xu et al., 2015; Zhang et al., 2003)), and heterogeneous directional propagation patterns that extended beyond those seen previously in humans.

Our findings show that the human cortex exhibits complex spatial patterns during memory processing that are visible through the propagation of oscillations. By demonstrating rotational, concentric, and other traveling wave patterns that were specific to particular functional states, our findings are consistent with predictions from computational models suggesting that traveling waves would exhibit individualized patterns based on different cognitive and computational processes, as well as variations in anatomical connectivity (Bhattacharya et al., 2021; Sato, 2022).

## Results

### Theta/alpha and beta frequency oscillations are widespread and organized as spatiotemporally stable traveling waves

To probe the role of brain oscillations and traveling waves in organizing the spatial and temporal structure of cortical activity during memory, we examined human electrocorticographic (ECoG) recordings from surgical epilepsy patients as they performed spatial and verbal memory tasks (**Methods, Figure 1, Supplementary Figure 1**). We sought to identify brain oscillations that were spatially organized into traveling waves and test whether they reorganize into different directional patterns to distinguish cognitive states. To examine this hypothesis, we developed a flexible framework for measuring traveling waves in ECoG recordings, quantifying their instantaneous spatial structure, and identifying spatial patterns that differentiate individual cognitive states.

**Figure 1:**
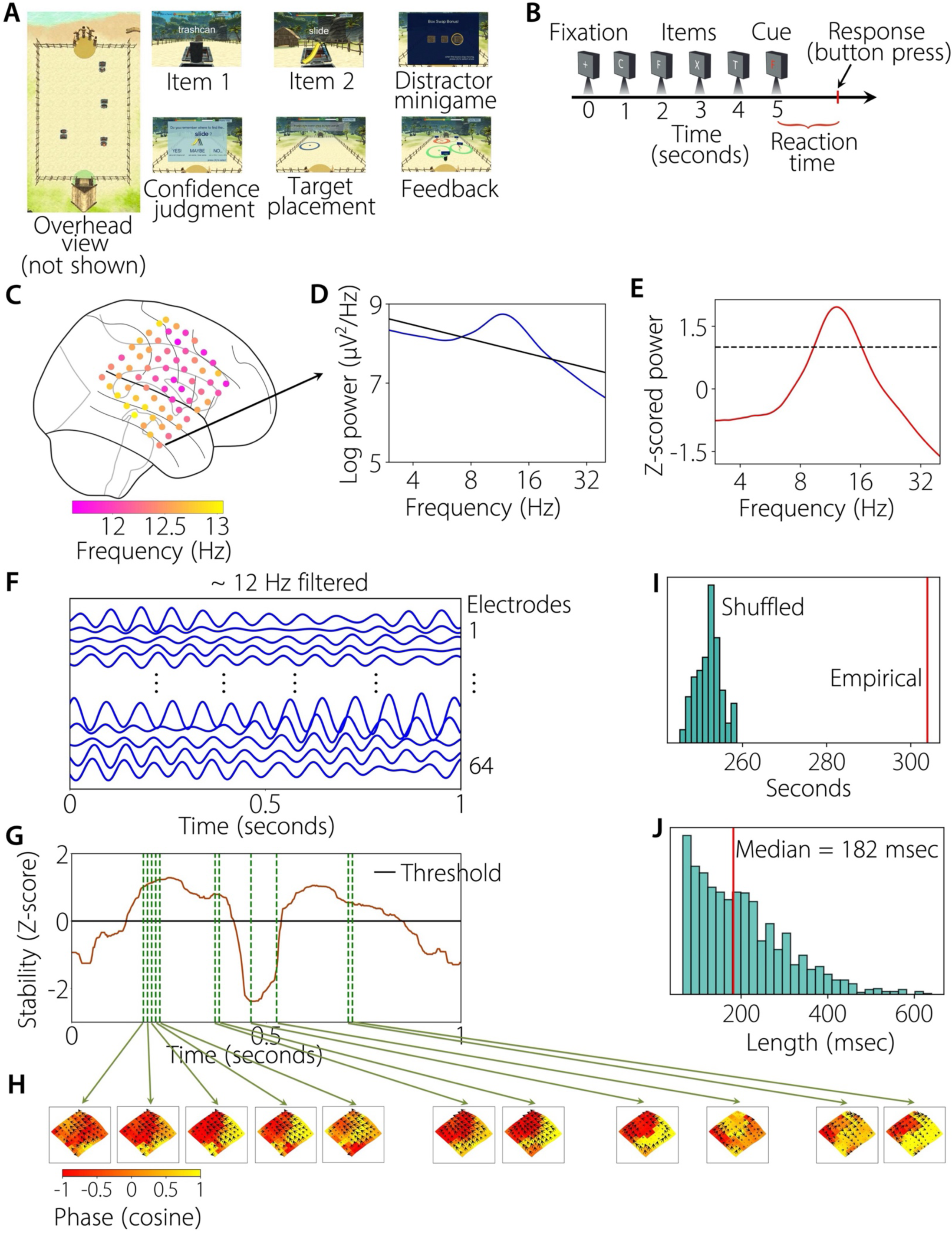
Task structure and identification of stable periods of traveling waves. **(A) Spatial episodic memory task.** Patients #1-9 (**Supplementary Figure 1**) performed multiple trials of a Treasure Hunt spatial memory task where they navigated to treasure chests located in a virtual environment containing various objects and after a short delay, were asked to retrieve the spatial location of the objects (**Methods**). **(B) Sternberg verbal working memory task.** Patients #10-24 (**Supplementary Figure 1**) performed multiple trials of a Sternberg verbal working memory task where they were presented with a list of English letters to silently hold the identity of each item in memory and during the response period, indicated whether the probe was present in the just-presented list (**Methods**). **(C-E) Identification of narrowband oscillation frequency.** The first step in detecting traveling waves is to identify the narrowband oscillation frequencies of electrodes. Shown in **C** are narrowband oscillation frequencies of each electrode in the ECoG grid of patient #21. We used Morlet wavelets to compute the power of the neural oscillations at 200 frequencies logarithmically spaced from 3 to 40 Hz (blue line in **D**). To identify narrowband oscillations at each electrode, we fit a line to each patient’s mean power spectrum in log–log coordinates using robust linear regression (Das et al., 2022) (black line in **D**). We then subtracted the actual power spectrum from the regression line. This normalized power spectrum (red line in **E**) removes the 1/f background signal and emphasizes narrowband oscillations as positive deflections (**E**). We identified narrowband peaks in the normalized power spectrum as any local maximum greater than one standard deviation above the mean (dotted black line in **E**). **(F) Filtered signals.** We filtered the signals of the electrodes in each cluster at their peak frequencies. Shown here are the filtered signals from an example trial of electrodes 1-64 in the ECoG grid shown in **C**. **(G, H) Traveling waves and identification of stable epochs.** Visual observation of the propagation of the absolute phase (Hilbert-transformed phase) across time showed the presence of traveling waves, here **H** shows a sink traveling wave corresponding to the electrode grid in **C** (arrows denote the direction of the wave, lengths of the arrows denote wave strength, and colors denote the cosine of the phase). We used a localized circular-linear regression approach to estimate traveling waves in each patient individually (Das et al., 2022) (**Methods**). We then identified stable periods of wave propagation (**Methods**). Shown in **G** are the stability values for an example trial from patient #21. Black line in **G** denotes the stability threshold. In the example trial shown here, there were two stable epochs. Dotted vertical green lines correspond to time-points for which example traveling waves are shown in **H**. The traveling waves operated in the stable regime for a few tens of milliseconds, then they entered into the unstable regime where the stable wave pattern broke down and a new wave pattern emerged, and then finally moving onto a new stable regime (Compare traveling waves in **H** corresponding to the dotted green lines within a stable epoch versus waves in **H** corresponding to dotted green lines in the unstable epoch or another stable epoch). **(I) We additionally used shuffling procedures as control which suggested that the observed stable epochs are not due to chance (Methods). (J) Distribution of the length of the stable epochs across trials for patient #21.**

To flexibly detect spatial patterns of traveling waves in each subject, we first identified the frequencies where groups of contiguous electrodes showed common oscillations (**Methods, Figures 1C–E**). Identifying a single common frequency is crucial because, by definition, a traveling wave involves an oscillation at a single frequency that progressively propagates across a region of cortex, thus making it possible to detect the traveling wave when it passes by these electrodes (Das et al., 2022). Overall, oscillations were most often present in the theta, alpha, and beta frequency bands (**Figure 1C, Supplementary Tables 1, 2**), and ∼86% of all electrodes showed a narrowband oscillation in at least one of these bands. Interestingly, sometimes the same ECoG grid showed multiple peaks in the power spectrum, i.e., they had peaks at both the theta/alpha and beta frequency bands (**Supplementary Tables 1, 2**, also see **Figure 8**), which suggests the presence of multiple overlapping oscillatory networks.

We next distinguished the patterns of oscillations that were traveling waves. Because a traveling wave involves an oscillation at a single frequency that is present simultaneously on multiple electrodes, we identified the contiguous clusters of electrodes with oscillations at the same frequency. A traveling wave requires the presence of a systematic propagation of phase across a cluster of electrodes. To test for this pattern, we filtered the signals of the electrodes in each cluster at their peak frequencies and extracted the instantaneous phase at each electrode and timepoint using the Hilbert transform (**Figure 1F**). Visual observation of the propagation of the absolute phase (Hilbert-transformed phase) across time showed clear traveling waves, with phase advancing in consistent spatial patterns across neighboring electrodes (see **Figure 1H**, for example).

Next, to probe this phenomenon systematically, we implemented a quantitative framework for measuring the spatial pattern of each traveling wave. Here, for each electrode in a cluster, we used a windowed circular–linear regression approach to identify the relative phases that exhibit a linear relationship with electrode locations *locally* (Das et al., 2022) (**Methods**). This regression also measured the properties of each wave’s propagation, including its phase velocity and direction at each timepoint (**Figure 1H, Methods**).

Recent evidence has shown that oscillations and traveling waves in humans often appear in transient bursts, but that they nonetheless are functionally relevant (Freeman & Rogers, 2002; Muller et al., 2018; Roberts et al., 2019; Schmidt et al., 2023; van Ede et al., 2018; van Vugt et al., 2007). We thus designed a single-trial method that identified stable periods of wave propagation (**Methods, Figure 1G**) and used this procedure to identify individual epochs where waves progressed across the cortex in a consistent spatial arrangement (**Methods, Figures 1G, H, Supplementary Videos 1-5**). This procedure revealed that the stable periods of individual traveling waves lasted ∼80–180 msec on average (**Figures 1G, H**). These wave epochs were statistically robust (all *p’s* < 0.001, **Methods, Figures 1I, J, Supplementary Figure 2**) ruling out the possibility that these stable epochs could arise due to chance or noise, and their period of stability was shorter for oscillations at high frequencies than low (median 107 versus 134 msec, *p* = 0.002, Mann-Whitney U-test). Consistent with earlier work (Das et al., 2022; Halgren et al., 2019; Stolk et al., 2019; Zhang et al., 2018), the phase velocities of these traveling waves was ∼0.3-3.3 m/s, and faster for oscillations at higher temporal frequencies than low (median 1.1 m/s for 12-26 Hz versus 0.5 m/s at 5-12 Hz, *p* < 0.001, Mann-Whitney U-test; (Zhang et al., 2018)).

### Independent component analysis (ICA) reveals the most dominant traveling wave patterns

Using this approach, we observed a diverse range of robust traveling wave patterns across subjects (**Figures 2, 3**). These patterns included plane waves as seen previously, but also spirals, concentric waves, and spatially heterogeneous, complex waves (for examples, see **Supplementary Figures 7-16**). We also saw that over time the traveling waves at individual electrodes often shifted between different patterns. To quantitatively distinguish the full diversity of spatial wave patterns over time, we used an algorithm based on independent component analysis (ICA) (Fu et al., 2015; Li & Adalı, 2010) to identify the range of spatial patterns of traveling waves that appeared on each electrode cluster (see **Methods**). Using this algorithm, we labeled each category of spatial wave pattern that appeared on each cluster, which we refer to as a “mode” (**Figure 2, Methods**). We classified each mode according to its shape, including plane, spiral, and concentric waves, as well as other waves with complex shapes (**Figure 3**). This algorithm provided a series of weights that quantify the contribution of each mode to the epoch’s current shape. We call these weights the epoch’s “activation function”, and they quantify the instantaneous magnitude and direction of a mode for each epoch. Thus, by examining the modes and activation functions, it shows quantitatively the types of traveling wave patterns that were most strongly present at each moment in the recording, with the magnitude of the activation function indicating how strongly each mode was represented.

**Figure 2:**
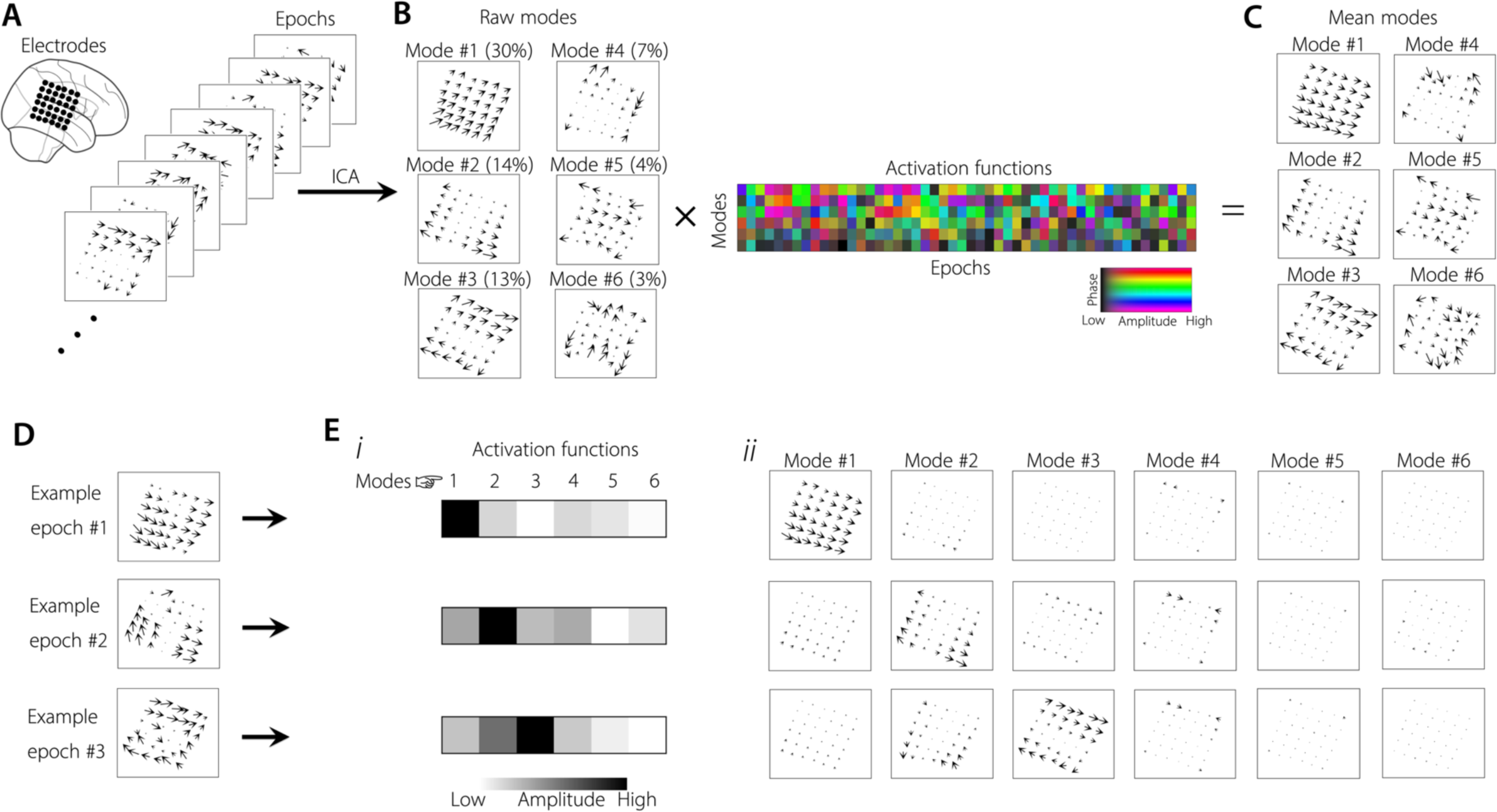
Independent component analysis (ICA) of traveling waves. **(A) Wave patterns across stable epochs are concatenated and passed as input to ICA (Methods).** Shown are example stable epochs from patient #5 (∼6.1 Hz traveling wave). **(B) Raw modes. We extracted the independent activation functions (or, weights) and the corresponding modes (“raw modes”) as the output from the ICA (Methods).** Variance explained by each mode is shown in brackets. Activation functions are complex and shown in color with colorbar. **(C) Mean modes. Multiplication of each of the raw mode with the mean of the weights across all stable epochs corresponding to that specific mode results in a unique wave pattern (“mean modes”) associated with that mode. (D-E) ICA modes are present at the individual epoch level.** Each individual epoch can be represented as the sum of the products of activation functions and modes. At the individual epoch level, a higher ICA weight for that epoch corresponding to a specific mode indicates higher representation of that wave pattern in that specific epoch and a lower ICA weight for an epoch corresponding to a specific mode indicates lower representation of that wave pattern in that specific epoch (**E**). Shown are three example epochs (**D**), with the corresponding activation functions for the top 6 modes (**Ei**), and the corresponding modes (**Eii**). Note that in **Eii**, the modes are weighted by the corresponding activation functions, with arrows denoting the wave strength.

**Figure 3:**
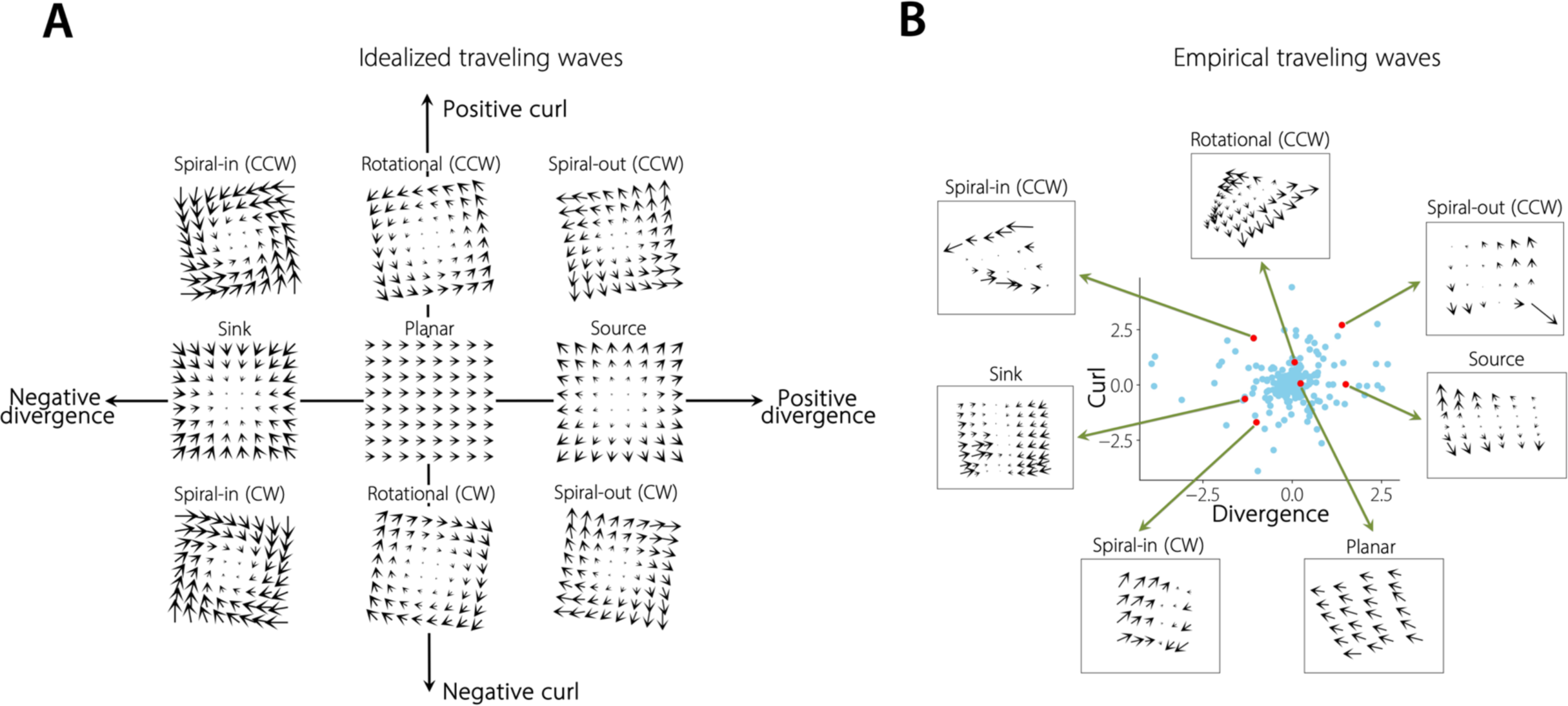
Classification of traveling waves. **(A) Simulated examples (idealized traveling waves) of planar, rotational (clockwise/counter-clockwise), and concentric (source/sink) waves, in the divergence-curl plane. (B) Experimental data (empirical traveling waves) shown as data points (blue dots), along with the associated wave patterns, in the divergence-curl plane.** After extracting the mean modes from the ICA, we next classified each of the mean modes as one of “planar”, “rotational”, “concentric” (“expanding” or “contracting”), or “complex” categories (**Methods**).

This ICA algorithm effectively quantified the spatial wave patterns that appeared visually in the recordings, as the dominant wave patterns that were evident from visual inspection generally corresponded with the modes with the strongest weights from ICA. For example, **Figure 2D** shows three different wave patterns seen in the raw data, which match the corresponding activation functions for the top 6 modes shown in **Figure 2Ei**, and the corresponding modes shown in **Figure 2Eii** (also see **Supplementary Figures 5, 6**).

Using this ICA procedure to identify the modes that were present on each cluster throughout the task, we identified a wide range of spatial traveling wave patterns across all clusters (**Methods, Figure 2, Supplementary Figures 7-16**). Individual clusters showed a mean of 6 significant modes, indicating that there were generally multiple spatial patterns of traveling waves within individual clusters. On average the first three modes explained ∼34%, ∼19%, and ∼10% variance respectively, and the first six modes combined explained >80% variance (**Figure 4E**). In this way, using ICA, we can identify the diverse traveling wave patterns that are present at each oscillation cluster, and reveal how waves with different shapes vary throughout the task.

**Figure 4:**
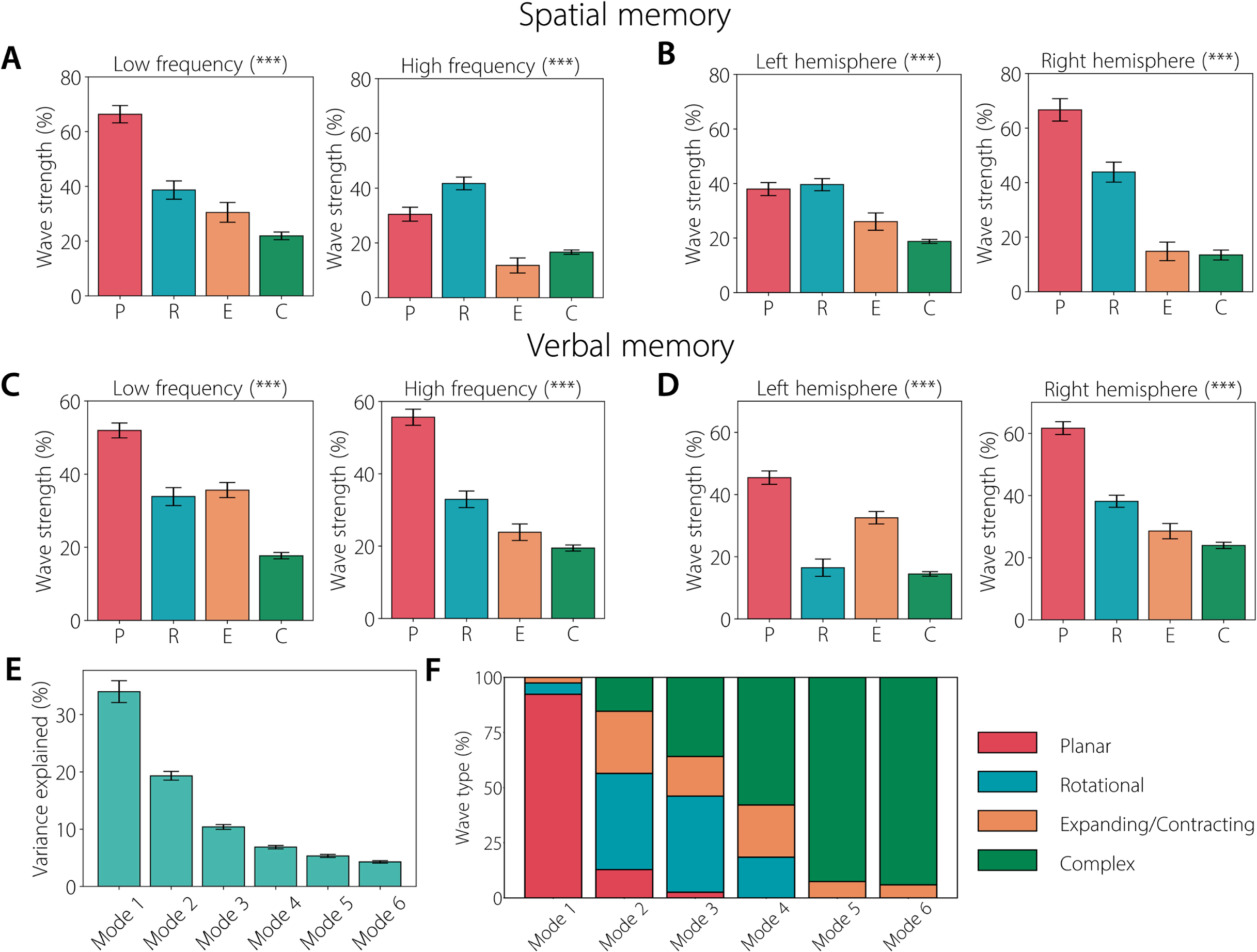
Population-level characteristics of different wave patterns. **(A) Wave strength for each category of wave (P: Planar, R: Rotational, E: Expanding/Contracting, C: Complex) for low (left panel) and high (right panel) frequency in the spatial memory task.** Error bars denote standard error of the proportion across modes. **(B) Wave strength for each category of wave for left (left panel) and right (right panel) hemisphere in the spatial memory task. (C) Wave strength for each category of wave for low (left panel) and high (right panel) frequency in the verbal memory task. (D) Wave strength for each category of wave for left (left panel) and right (right panel) hemisphere in the verbal memory task. (E) Variance explained for each mode across all clusters and tasks.** Error bars denote standard error of the proportion across clusters. **(F) Percentage of each wave type in each mode.** *** *p* < 0.001 (FDR-corrected).

### Diverse spatial patterns of traveling waves and their characteristics

To explain the diversity of the spatial wave patterns in the data, we classified each identified mode based on their shape into one of the following categories: “planar”, “rotational”, “concentric” (“expanding” or “contracting”), or “complex” (**Methods, Figure 3**) (Townsend et al., 2015). Complex waves were those that showed a robust spatial propagation pattern (i.e., *p’s* < 0.001) but did not meet the criteria for the other categories. Some complex waves showed a combination of multiple patterns, such as separate subsets of electrodes showed planar, rotating, or expanding/contracting waves (**Figure 9, Supplementary Figures 7-16**).

One previous study showed rotational and expanding traveling waves during human sleep spindles, supporting the idea that these types of waves may support memory consolidation and plasticity (Muller et al., 2016). Based on this, we hypothesized that rotational and expanding (sources) traveling waves would be more prevalent in the spatial episodic memory task which involves processing a broad range of inputs including spatial and non-spatial information.

The spatial organization of traveling waves varied across regions and temporal frequencies. For clusters with oscillations at low temporal frequencies, plane waves were most dominant (median wave strength 59% across tasks), and this pattern was present in both the spatial (*ξ*^2^(3) = 165.0, *p* < 0.001, **Figure 4A**) and verbal (*ξ*^2^(3) = 288.7, *p* < 0.001, **Figure 4C**) memory tasks. At higher frequencies, although plane waves were again most dominant overall (median wave strength 43% across tasks), there was an increased prevalence of rotational waves (median wave strength 37% across tasks), especially in the spatial memory task (*ξ*^2^(3) = 162.3, *p* < 0.001, **Figure 4A**). When we compared the strength of these different traveling wave patterns as indexed by variance explained (see **Methods**), we found that planar waves were strongest, followed by rotating and expanding/contracting waves (**Figures 4E, F**). Complex waves, although highly statistically significant and strongly relevant to behavior (see below), were less strong and explained less variance in the raw data compared to the other wave types.

We also compared these effects between hemispheres, in light of work on lateralized oscillations in hippocampus and neocortex (Das et al., 2022; Miller et al., 2018). Although plane waves were dominant in both hemispheres, they were stronger on the right hemisphere (∼64%) than the left (∼42%; *χ*^2^ test *p* < 0.01, **Figures 4 B, D**). We next compared this lateralization between the two different memory tasks. In the left hemisphere, during the spatial memory task wave patterns were preferentially rotational rather than concentric, (*χ*^2^(3) = 150.0, *p* < 0.001, **Figure 4B**). Conversely, concentric wave patterns were more common in the verbal memory task (*χ*^2^(3) = 288.1, *p* < 0.001, **Figure 4D**). But, in the right hemisphere, we found statistically similar patterns across both tasks. This suggests that the shape of traveling waves across the cortex during behavior reveal specific task-related neural assemblies that are lateralized across the cortex.

Given that we saw both counter-clockwise and clockwise rotational wave patterns, we examined whether the orientation of wave propagation was functionally relevant. One previous study showed that during sleep, human cortical spindles are rotational traveling waves that propagated in a temporal→parietal→frontal (TPF) direction (Muller et al., 2016). Therefore, we examined whether rotating waves in our dataset had a directional preference by labeling each one’s direction (TPF or TFP). If there were multiple rotational modes in a single oscillation cluster, then we labelled each separately. We did not find a significant preference for a specific rotational direction in either task. In the spatial memory task, there were 7 TPF waves versus 9 TFP waves (*p* > 0.05, binomial test) and, similarly, in the verbal memory task (12 TPF versus 15 TFP, *p* > 0.05, binomial test; **Supplementary Table 3**).

We also compared the propagation of concentric traveling waves, comparing the prevalence of inward (sink) versus outward (source) propagation. In the spatial task, there were more sources compared to sinks, as 78% (7 of 9) concentric waves were sources. Inversely, in the verbal memory task, there were more sinks than sources, as 68% (15 of 22) concentric waves were sinks (**Supplementary Table 3**). Thus, the brain shows different types of concentric waves (*χ*^2^ test, *p* < 0.02) between spatial and verbal memory.

### Traveling waves can distinguish both broad and specific cognitive states of human memory representations

Next, to identify the potential functional role of traveling waves, we examined how the prevalence of traveling waves with different shapes shifted between stages of memory. First, we examined encoding, retrieval, navigation, etc. in the spatial memory task. To assess statistical significance, we used multivariate analysis of variance (MANOVA) (**Methods**) to test for changes in activation functions between conditions.

Traveling waves showed different propagation patterns between stages of memory processing (**Figure 5, Supplementary Figures 7-16, Supplementary Videos 1-2**). As an example, **Figures 5A-C** show a cluster of electrodes that showed three distinct modes of traveling waves that changed their propagation direction between the stages of memory. In this cluster, the plane wave pattern (mode 1) was nearly absent during navigation and distractor stages (as indicated by its low magnitude) but was strongly present in other task phases. However, its direction of propagation shifted between behaviors, with a different direction for retrieval compared to the encoding, confidence, and feedback periods. Similarly, this cluster’s mode 2 also differed in propagation direction between memory stages and had the highest wave strength during retrieval and feedback. These state-specific traveling waves were prominent across our dataset, as all 13 oscillation clusters showed significant shifts in traveling wave direction or strength between stages of the spatial memory task.

**Figure 5:**
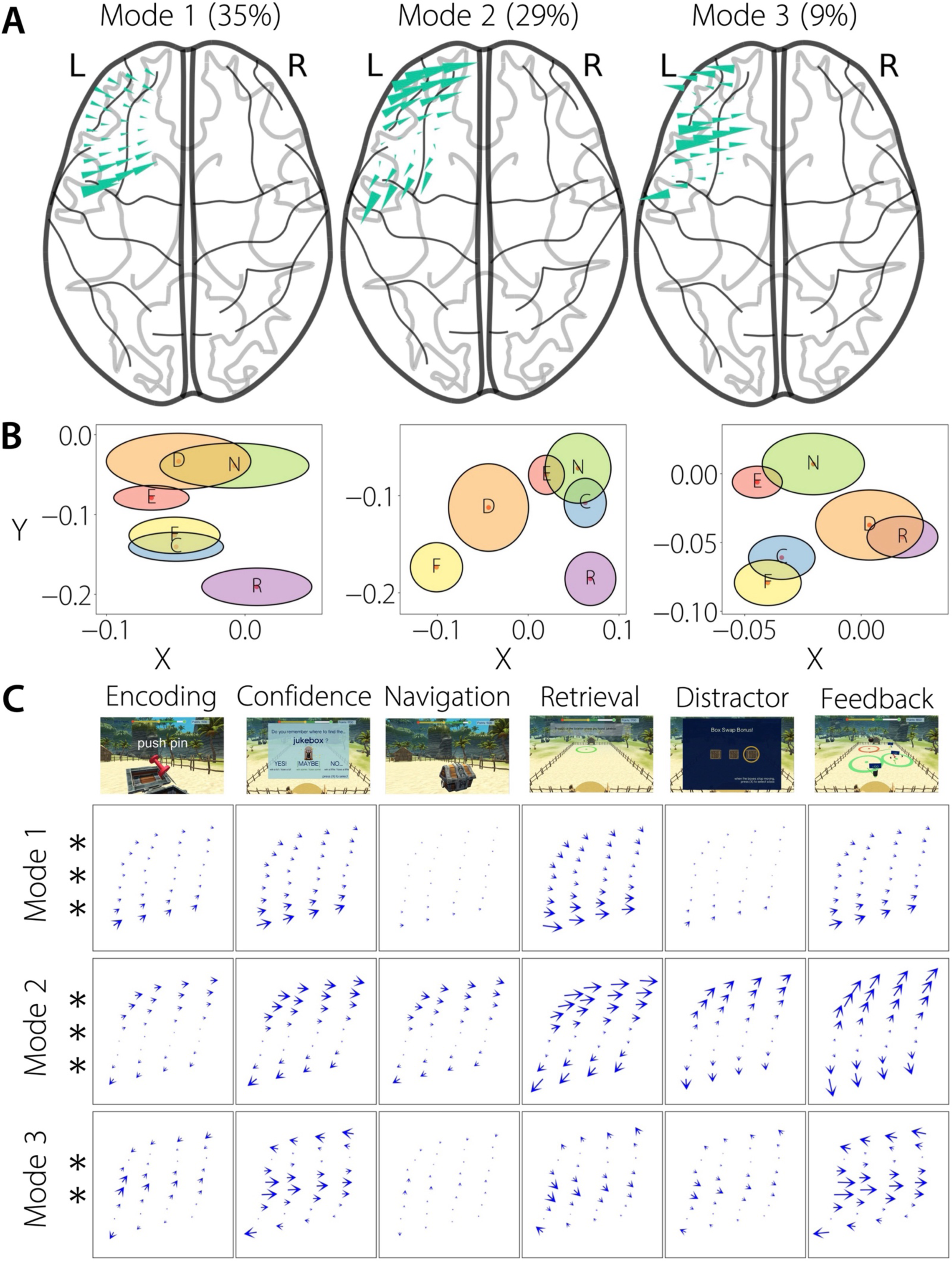
Traveling waves can distinguish broad cognitive states in human memory processing. **(A) Top 3 mean modes of patient #3 (∼17 Hz traveling wave) visualized on a brain surface plot.** Variance explained by each mode is indicated in brackets. **(B) Distinguishing cognitive states in this patient in the spatial memory experiment, shown are the activation functions in the complex plane for the three modes in A.** The shift in direction and/or strength of the traveling waves between different behaviors can be visualized in terms of the activation functions where, a change in the direction of the waves corresponds to a change in the angle of the activation functions (for example, compare confidence vs. distractor for mode 2 in **B and C**), a change in the strength of the waves corresponds to a change in the magnitude/length of the activation functions (for example, compare confidence vs. navigation for mode 1 in **B and C**), a change in both the direction and strength of the waves corresponds to a change in both the angle and length of the activation functions (for example, compare navigation vs. retrieval for mode 3 in **B and C**). For each ellipse (task period), the major axis (horizontal axis) denotes the standard-error-of-the-mean (SEM) for the real-part and the minor axis (vertical axis) denotes the SEM for the imaginary part, of the activation functions. E: Encoding, C: Confidence, N: Navigation, R: Retrieval, D: Distractor, F: Feedback. **(C) Mean modes for each cognitive state for the three modes in (A) in this patient.** Traveling waves changed either their direction (for example, confidence vs. distractor in mode 2), strength (for example, confidence vs. navigation in mode 1), or both (for example, navigation vs. retrieval in mode 3), to distinguish broad cognitive states in the spatial memory task. *** *p* < 0.001, ** *p* < 0.01 (FDR-corrected).

We also examined whether the properties of traveling waves shifted to represent the specific item that each subject was viewing, extending item-specific gamma oscillations seen previously (Jacobs & Kahana, 2009). In the verbal memory task, we found that many clusters showed traveling waves with different propagation directions and strengths for the encoding of individual items. For example, **Figures 6A-C** show a cluster of electrodes with planar traveling waves (on mode 1) that propagated consistently posteriorly when the subject viewed any letter except for “G”. This same cluster also showed a direction-shifting traveling wave in mode 3 which propagated in an antero-superior direction for letters “G” and “Q” and the opposite direction for letters “D” and “J”. These item-specific wave shifts were common, as 13 of 26 clusters in this task had traveling waves that shifted direction or strength for individual letters. Thus traveling waves often distinguish both broad and specific cognitive states of human memory representations (**Supplementary Figures 7-16**).

**Figure 6:**
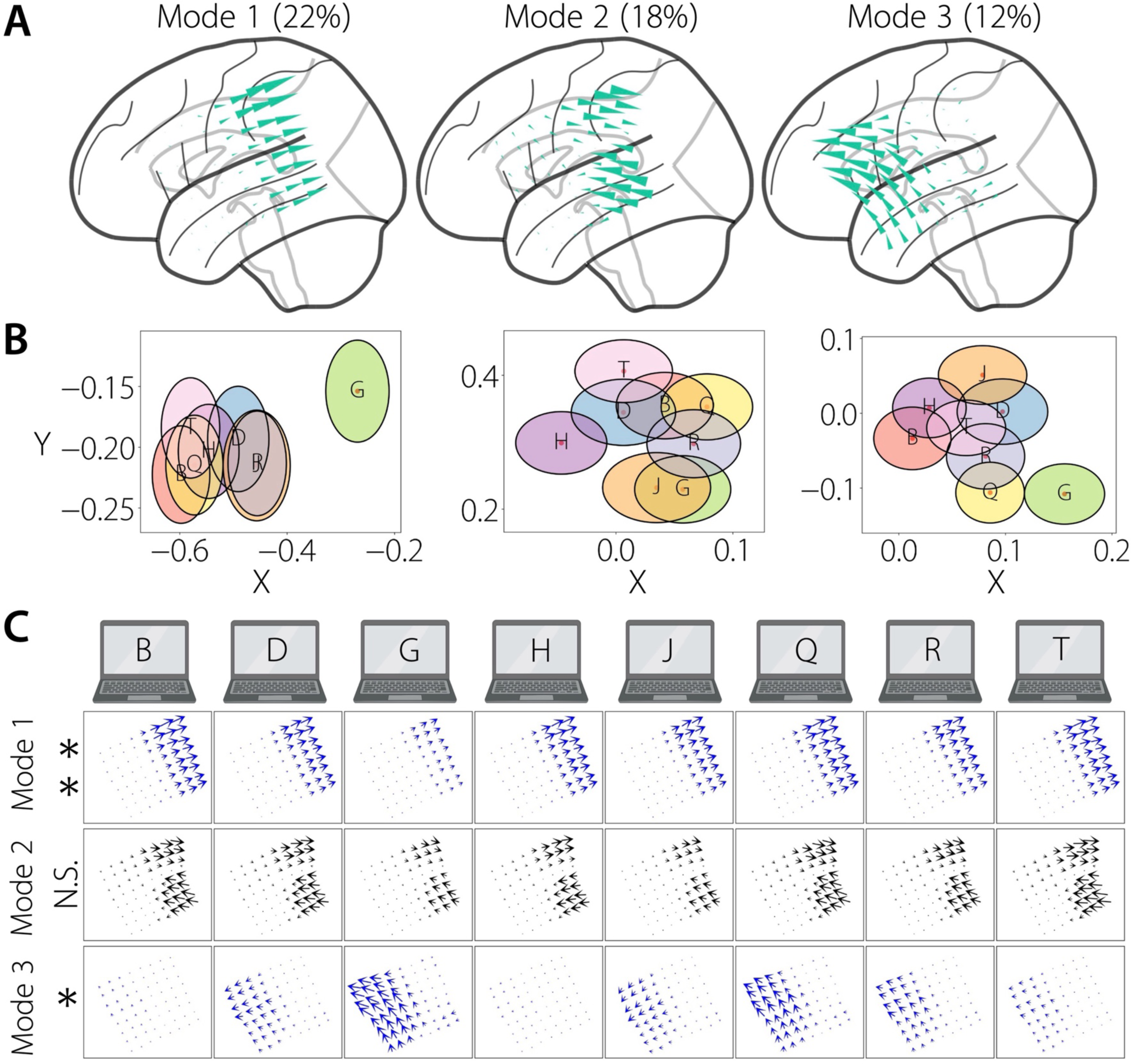
Traveling waves can distinguish specific cognitive states in human memory processing. **(A) Top 3 mean modes of patient #23 (∼7 Hz traveling wave) visualized on a brain surface plot.** Variance explained by each mode is indicated in brackets. **(B) Distinguishing specific cognitive states in this patient, shown are the activation functions in the complex plane for the three modes in A.** Similar to the spatial memory task, the shift in direction and/or strength of the traveling waves for the English letters can also be visualized in terms of the activation functions in the complex plane. **(C) Mean modes for each letter for the three modes in A in this patient.** Similar to the spatial memory task, traveling waves changed either their direction (for example, “J” vs. “R” in mode 3, also see **B**), strength (for example, “D” vs. “G” in mode 1, also see **B**), or both (for example, “G” vs. “H” in mode 3, also see **B**), to distinguish specific cognitive states in the verbal memory task. ** *p* < 0.01, * *p* < 0.05 (FDR-corrected).

### Waves that distinguish cognitive states are of diverse patterns and widespread across the human brain

An important finding of our work is showing that rotational, concentric, and complex patterns of waves also distinguish cognitive states, in addition to planar waves. Therefore, our next objective was to quantify the behavioral relevance of these non-planar traveling waves.

Similar to plane waves, rotational waves were also stable at the individual epoch level, distinguished both broad and specific cognitive states, and were widespread across the frontal, temporal, and parietal lobes (**Figures 7A-D, Supplementary Video 3, Supplementary Figures 7-16**). These patterns were evident in both the spatial and verbal memory tasks (**Figures 7E, G**, also see **Supplementary Figures 7-16**). For rotational waves, the wave strength was higher during the distractor period compared to the other task periods, especially in the left hemisphere (*χ*^2^(5) = 67.0, *p’s* < 0.001, **Figure 7F**).

**Figure 7:**
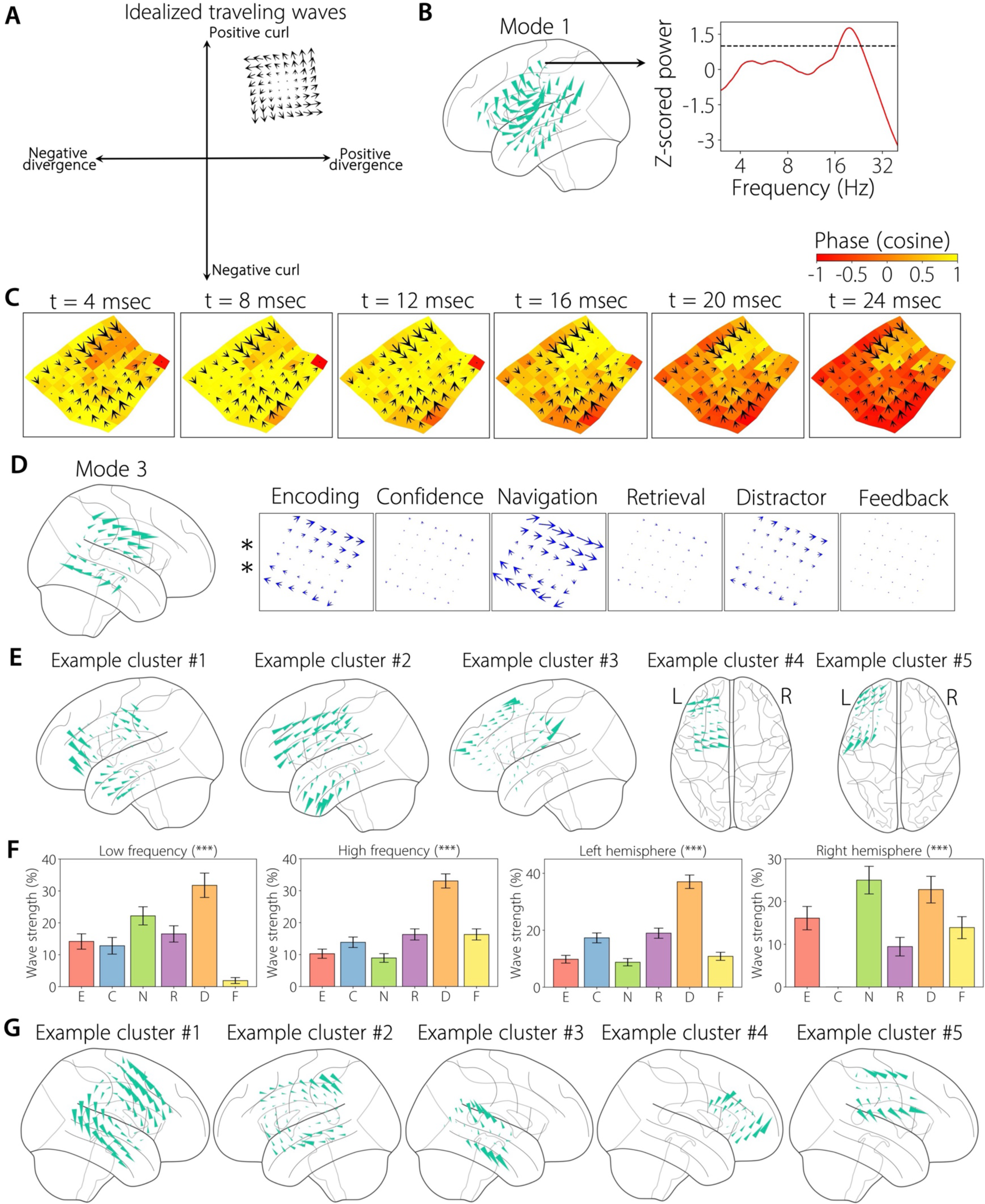
Rotational traveling waves can distinguish cognitive states in human memory processing. **(A) Simulated example of a rotational traveling wave (counter-clockwise spiral-out) in the divergence-curl plane. (B) Left panel: Traveling waves (mean mode) visualized on a brain surface plot for patient #2, mode #1 (∼20.6 Hz traveling wave). Right panel: Normalized z-scored power from an example electrode. (C) Rotational traveling waves are stable at the individual epoch level.** Shown is the propagation of the traveling wave across time for an example stable epoch for this patient, with arrows denoting the traveling waves and colors denoting the cosine of the phases. **(D) Rotational traveling waves can distinguish cognitive states in the spatial memory task.** First panel: Traveling waves visualized on a brain surface plot for patient #5, mode #3 (∼6.1 Hz traveling wave). Panels 2-7: Mean traveling waves for different task periods. ** *p* < 0.01 (FDR-corrected). **(E) Examples of rotational traveling waves in the spatial memory task, visualized on a brain surface plot, demonstrating that rotational waves are widespread across multiple brain areas.** Panel 1: Traveling waves for patient #6, mode #2, cluster frequency (CF) ∼ 8.0 Hz. Panel 2: Traveling waves for patient #6, mode #2, CF ∼ 17.7 Hz. Panel 3: Traveling waves for patient #8, mode #2, CF ∼ 19.9 Hz. Panel 4: Traveling waves for patient #1, mode #2, CF ∼ 20.1 Hz. Panel 5: Traveling waves for patient #3, mode #2, CF ∼ 6.0 Hz. **(F) Wave strength (%) for each task period for rotational wave category in the spatial memory task, shown separately for low and high frequencies and left and right hemispheres.** E: Encoding, C: Confidence, N: Navigation, R: Retrieval, D: Distractor, F: Feedback. *** *p* < 0.001 (FDR-corrected). **(G) Examples of rotational traveling waves in the verbal memory task, visualized on a brain surface plot, demonstrating that rotational waves are widespread across multiple brain areas in the verbal memory task as well.** Panel 1: Traveling waves for patient #21, mode #2, CF ∼ 12.4 Hz. Panel 2: Traveling waves for patient #24, mode #3, CF ∼ 22.7 Hz. Panel 3: Traveling waves for patient #22, mode #4, CF ∼ 11.5 Hz. Panel 4: Traveling waves for patient #10, grid #1, mode #2, CF ∼ 17.5 Hz. Panel 5: Traveling waves for patient #16, mode #3, CF ∼ 9.2 Hz.

Concentric and complex wave patterns also distinguished both broad and specific cognitive states across brain regions and tasks (**Figures 8, 9, Supplementary Videos 4, 5**). Interestingly, during the distractor phase of the task, the strength of complex waves was significantly higher compared to other task periods across all frequencies and hemispheres (*χ*^2^(5) = 43.2, *p’s* < 0.001, **Figure 9F**), however, this pattern was not seen for the concentric waves (*χ*^2^(5) < 19.6, *p’s* > 0.01, **Figure 8F**).

**Figure 8:**
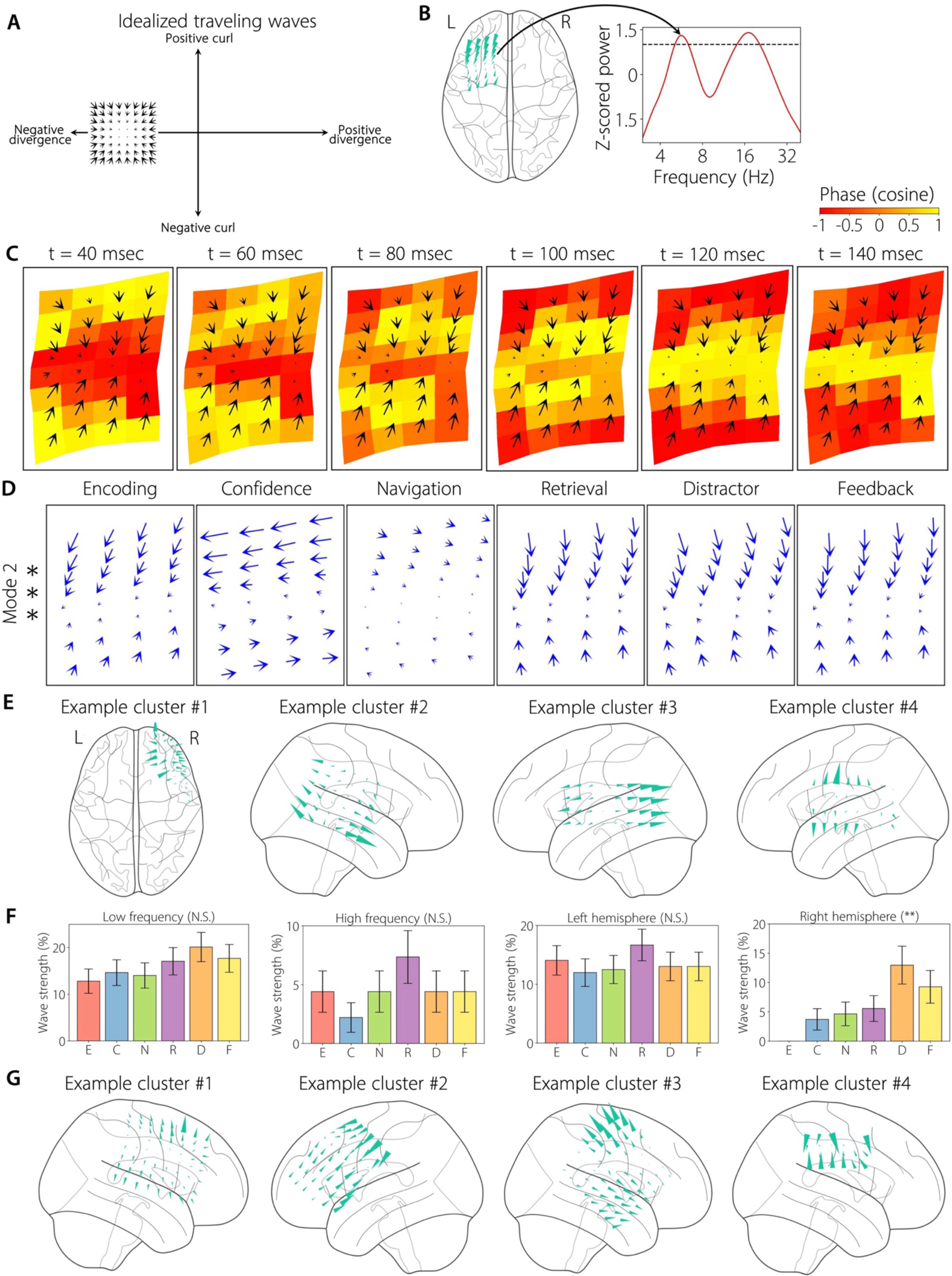
Concentric traveling waves can distinguish cognitive states in human memory processing. **(A) Simulated example of a sink in the divergence-curl plane. (B) Left panel: Traveling waves (mean mode) visualized on a brain surface plot for patient #1, mode #2 (∼5.6 Hz traveling wave). Right panel: Normalized z-scored power from an example electrode (note the presence of two peaks, one at lower frequency and another at higher frequency). (C) Concentric traveling waves are stable at the individual epoch level.** Shown is the propagation of the traveling wave across time for an example stable epoch for this patient. **(D) Concentric traveling waves can distinguish cognitive states in the spatial memory task.** Panels 1-6: Mean traveling waves for different task periods for this patient. *** *p* < 0.001 (FDR-corrected). **(E) Examples of concentric traveling waves in the spatial memory task, visualized on a brain surface plot, demonstrating that expanding/contracting waves are widespread across multiple brain areas.** Panel 1: Traveling waves for patient #4, mode #5, CF ∼ 22.5 Hz. Panel 2: Traveling waves for patient #5, mode #2, CF ∼ 6.1 Hz. Panel 3: Traveling waves for patient #9, mode #2, CF ∼ 9.7 Hz. Panel 4: Traveling waves for patient #9, mode #4, CF ∼ 9.7 Hz. **(F) Wave strength (%) for each task period for concentric wave category in the spatial memory task.** E: Encoding, C: Confidence, N: Navigation, R: Retrieval, D: Distractor, F: Feedback. ** *p* < 0.01, N.S. Not significant (FDR-corrected). **(G) Examples of concentric traveling waves in the verbal memory task, visualized on a brain surface plot, demonstrating that expanding/contracting waves are widespread across multiple brain areas in the verbal memory task as well.** Panel 1: Traveling waves for patient #11, mode #2, CF ∼ 19.8 Hz. Panel 2: Traveling waves for patient #19, mode #3, CF ∼ 6.5 Hz. Panel 3: Traveling waves for patient #21, mode #3, CF ∼ 25.4 Hz. Panel 4: Traveling waves for patient #16, mode #3, CF ∼ 20.0 Hz.

**Figure 9:**
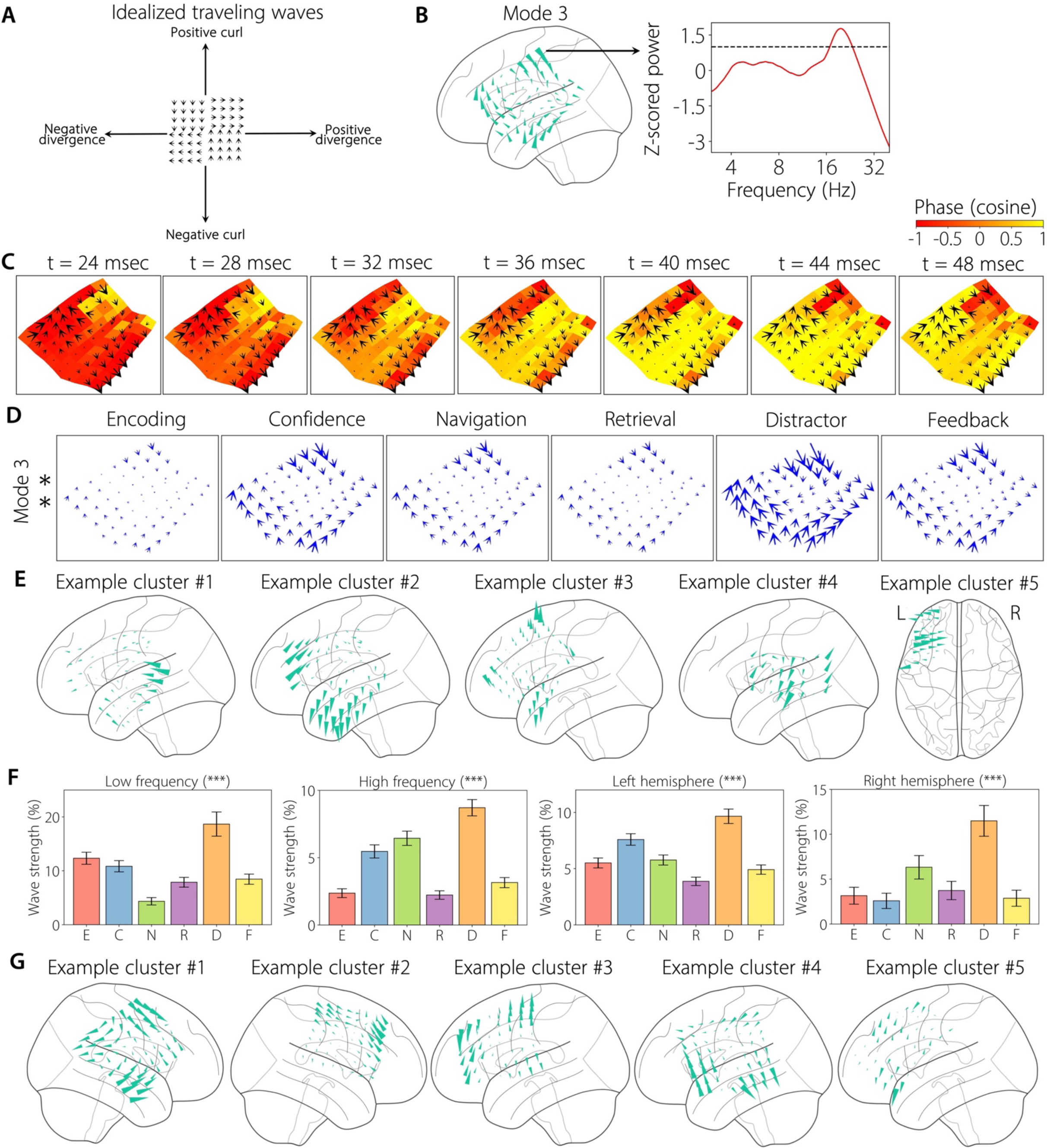
Complex patterns of traveling waves can distinguish cognitive states in human memory processing. **(A) Simulated example of a complex traveling wave in the divergence-curl plane. (B) Left panel: Traveling waves (mean mode) visualized on a brain surface plot for patient #2, mode #3 (∼20.6 Hz traveling wave). Right panel: Normalized z-scored power from an example electrode. (C) Complex traveling waves are stable at the individual epoch level.** Shown is the propagation of the traveling wave across time for an example stable epoch for this patient. **(D) Complex traveling waves can distinguish cognitive states in the spatial memory task.** Panels 1-6: Mean traveling waves for different task periods for this patient. ** *p* < 0.01 (FDR-corrected). **(E) Examples of complex traveling waves in the spatial memory task, visualized on a brain surface plot, demonstrating that complex waves are widespread across multiple brain areas.** Panel 1: Traveling waves for patient #6, mode #4, CF ∼ 17.7 Hz. Panel 2: Traveling waves for patient #6, mode #3, CF ∼ 8.0 Hz. Panel 3: Traveling waves for patient #8, mode #3, CF ∼ 19.9 Hz. Panel 4: Traveling waves for patient #9, mode #3, CF ∼ 9.7 Hz. Panel 5: Traveling waves for patient #3, mode #3, CF ∼ 17.2 Hz. **(F) Wave strength (%) for each task period for complex wave category in the spatial memory task.** E: Encoding, C: Confidence, N: Navigation, R: Retrieval, D: Distractor, F: Feedback. *** *p* < 0.001 (FDR-corrected). **(G) Examples of complex traveling waves in the verbal memory task, visualized on a brain surface plot, demonstrating that complex waves are widespread across multiple brain areas in the verbal memory task as well.** Panel 1: Traveling waves for patient #21, mode #3, CF ∼ 12.4 Hz. Panel 2: Traveling waves for patient #11, mode #3, CF ∼ 19.8 Hz. Panel 3: Traveling waves for patient #20, mode #4, CF ∼ 5.7 Hz. Panel 4: Traveling waves for patient #24, mode #2, CF ∼ 22.7 Hz. Panel 5: Traveling waves for patient #19, mode #5, CF ∼ 6.5 Hz.

Finally, we quantified the number of modes that were significant for each type of wave. The percentage of modes that were significant in each category of wave did not differ from each other, in any frequency or hemisphere (*χ*^2^(3) < 6.4, *p’s* > 0.05, **Figures 10A-D**). Moreover, the percentage of significance in each mode did not differ from each other (*χ*^2^(5) < 4.2, *p’s* > 0.05, **Figures 10E, F**).

**Figure 10:**
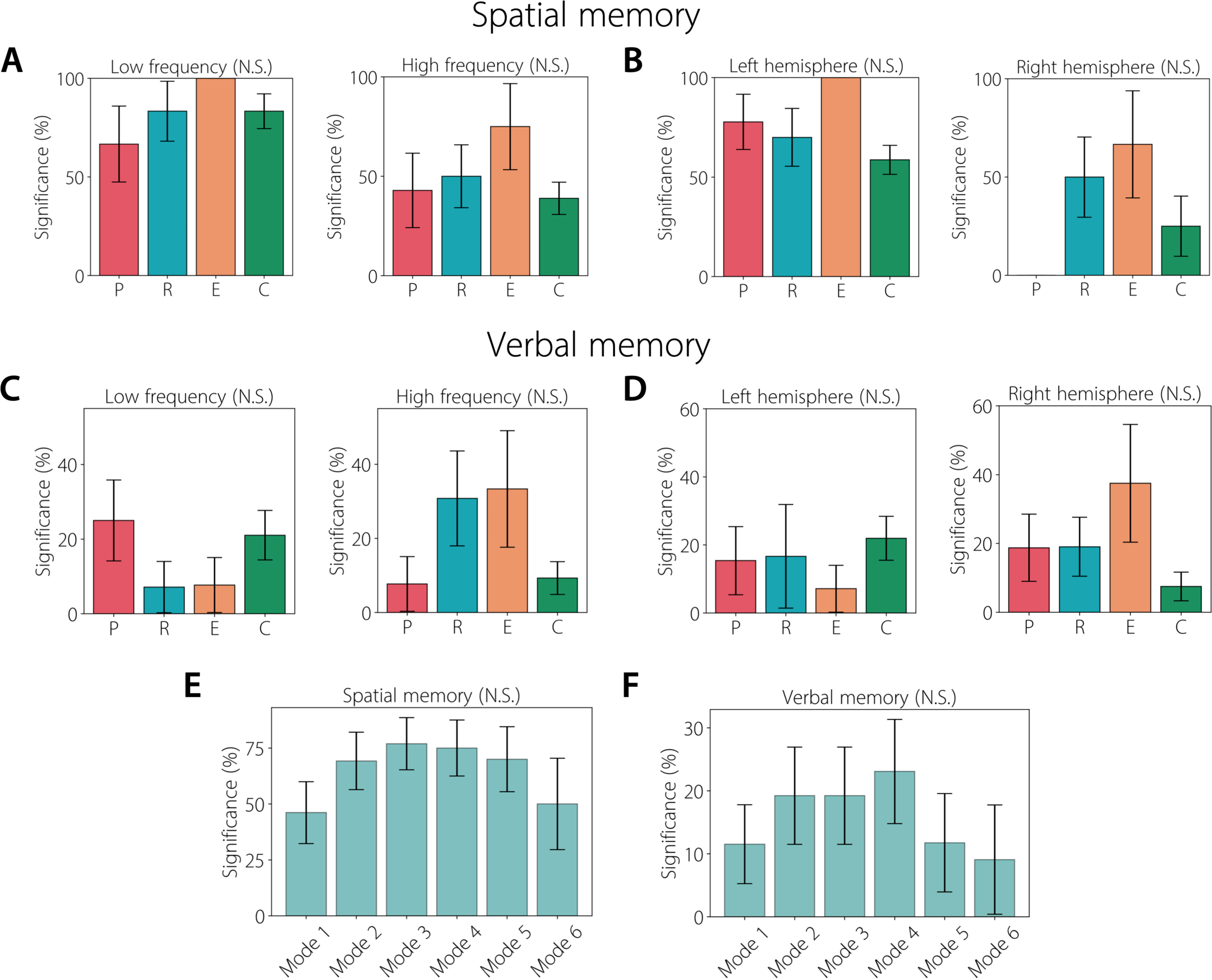
Population-level behavioral results. **(A) Percentage of significant modes for each category of wave (P: Planar, R: Rotational, E: Expanding/Contracting, C: Complex) for low (left panel) and high (right panel) frequency in the spatial memory task.** Error bars denote standard error of the proportion across modes. N.S. Not significant. **(B) Percentage of significant modes for each category of wave for left (left panel) and right (right panel) hemisphere in the spatial memory task. (C) Percentage of significant modes for each category of wave for low (left panel) and high (right panel) frequency in the verbal memory task. (D) Percentage of significant modes for each category of wave for left (left panel) and right (right panel) hemisphere in the verbal memory task. (E) Percentage of significance for each mode in the spatial memory task. (F) Percentage of significance for each mode in the verbal memory task.**

### Cognitive states of human memory representations can be decoded from traveling waves

Since the spatial patterns of traveling waves varied reliably with cognitive states, we hypothesized that we would be able to use these signals to support prediction and brain-computer interfacing. To decode each subject’s cognitive state from their spatial pattern of traveling waves, we trained multilayer neural networks, with cross-validation, for classifying pairwise cognitive states. We performed this decoding separately for the spatial and verbal memory tasks (**Methods, Figures 11, 12**). We used the extracted activation functions (weights) as inputs for the neural network classifiers.

**Figure 11:**
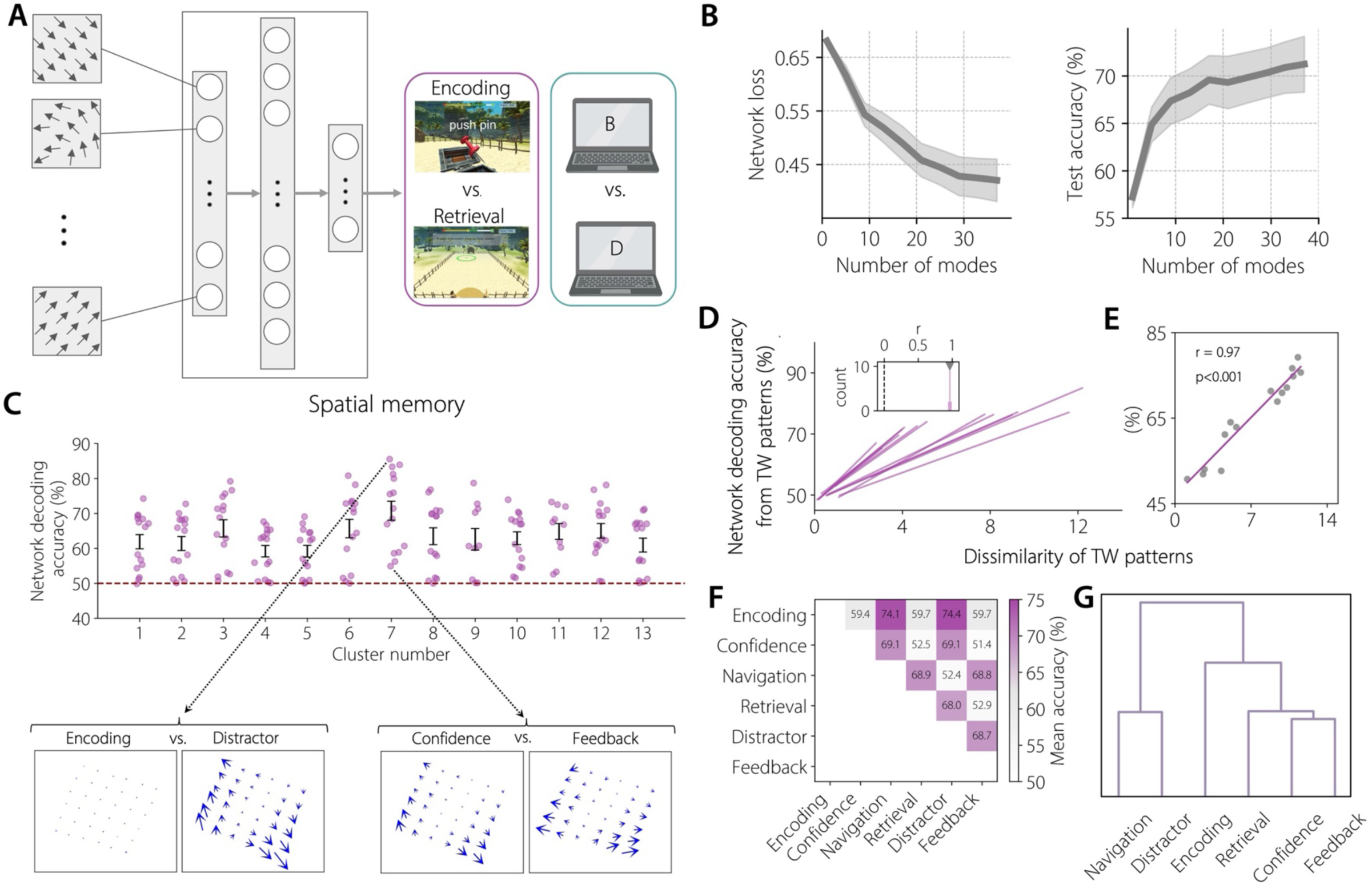
Decoding broad cognitive states of human memory representations from traveling waves. **(A) Neural network architecture.** We used a multilayer neural network with an input layer, two hidden layers (32 and 16 neurons respectively), and an output layer for decoding pairwise cognitive states (for example, encoding versus retrieval, letter “B” versus letter “D”, etc.) (see **Methods** for details). We used the extracted weights from the ICA procedure as features for training our neural network classifiers. **(B) Neural network loss (left panel) and test accuracy (right panel) versus number of modes.** Increasing the number of modes reduced the network training loss and increased test decoding accuracy. **(C) Network decoding accuracy for each oscillation cluster in the spatial memory task.** Each point in the plot corresponds to a pair of broad cognitive states (for example, encoding versus retrieval). Error bars show SEM decoding accuracies across all pairs. Dotted red line denotes chance level (50%). Inset shows that pairs of cognitive states which had higher decoding accuracy had wave patterns that were visually more distinct from each other. Example waves belong to patient #5, mode #2, CF ∼ 6.1 Hz. **(D, E) Network decoding accuracy versus dissimilarity of traveling wave patterns, in the spatial memory task.** Decoding accuracy was positively correlated (all *ps* < 0.001) with the dissimilarity of traveling wave (TW) patterns, demonstrating that the decoding results are consistent with the results obtained from the MANOVA analysis. One line was fit for each oscillation cluster and the inset shows the histogram of Pearson correlation values across clusters. **E** shows fitted line for an example oscillation cluster, with each point in the plot denoting a pair of broad cognitive states (for example, encoding versus retrieval). **(F) Mean decoding accuracy across all oscillation clusters for each pair of cognitive states, in the spatial memory task. (G) Dendrogram of F, demonstrating that the navigation and distractor states were the most decodable from the other states, but relatively less decodable from each other.**

We found that cognitive states can be reliably decoded at the individual cluster level and also at the group level (**Figure 11C**, *p* < 0.001, one-sided sign test), for the spatial memory task.

Interestingly, we found that some cognitive states can be more reliably decoded than others (**Figures 11C, F**). For example, **Figure 11C** shows that for cluster 7, the spatial wave patterns for the encoding and distractor periods were distinct from each other and hence were more easily distinguishable compared to those from the confidence and feedback periods. Overall, the navigation and distractor states were generally decodable from other states, but not from each other, which indicates that these behaviors generally exhibit similar traveling wave patterns (**Figures 11F, G**).

Similarly, we found that the identity of the specific memory item that a subject was viewing can be reliably decoded from the shape of their traveling waves in the verbal memory task, and also at the group level (**Figure 12A**, *p* < 0.001, one-sided sign tests across clusters versus chance). Overall, we often found that letter “H” was the most decodable, followed by the letters “J” and “Q”, compared to the other letters (**Figures 12D, E**). These results mean that letters “H”, “J”, and “Q” often elicited distinctive patterns of traveling waves (Jacobs & Kahana, 2009). Visually, cognitive states that showed lower decoding accuracy had more similar wave patterns compared to the ones that had higher decoding accuracy (**Figures 11C, 12A**), thus the results of this network decoding analysis were closely aligned with the results from the MANOVA analysis (**Figures 5, 6**). This similarity between the results of these analyses was robust, as we found mostly positive correlations between the neural network decoding accuracy and the dissimilarity of traveling wave patterns (calculated as the mean Euclidean distance between the ICA weights for the given pair of cognitive states) (**Figures 11D, E, 12B, C**).

**Figure 12:**
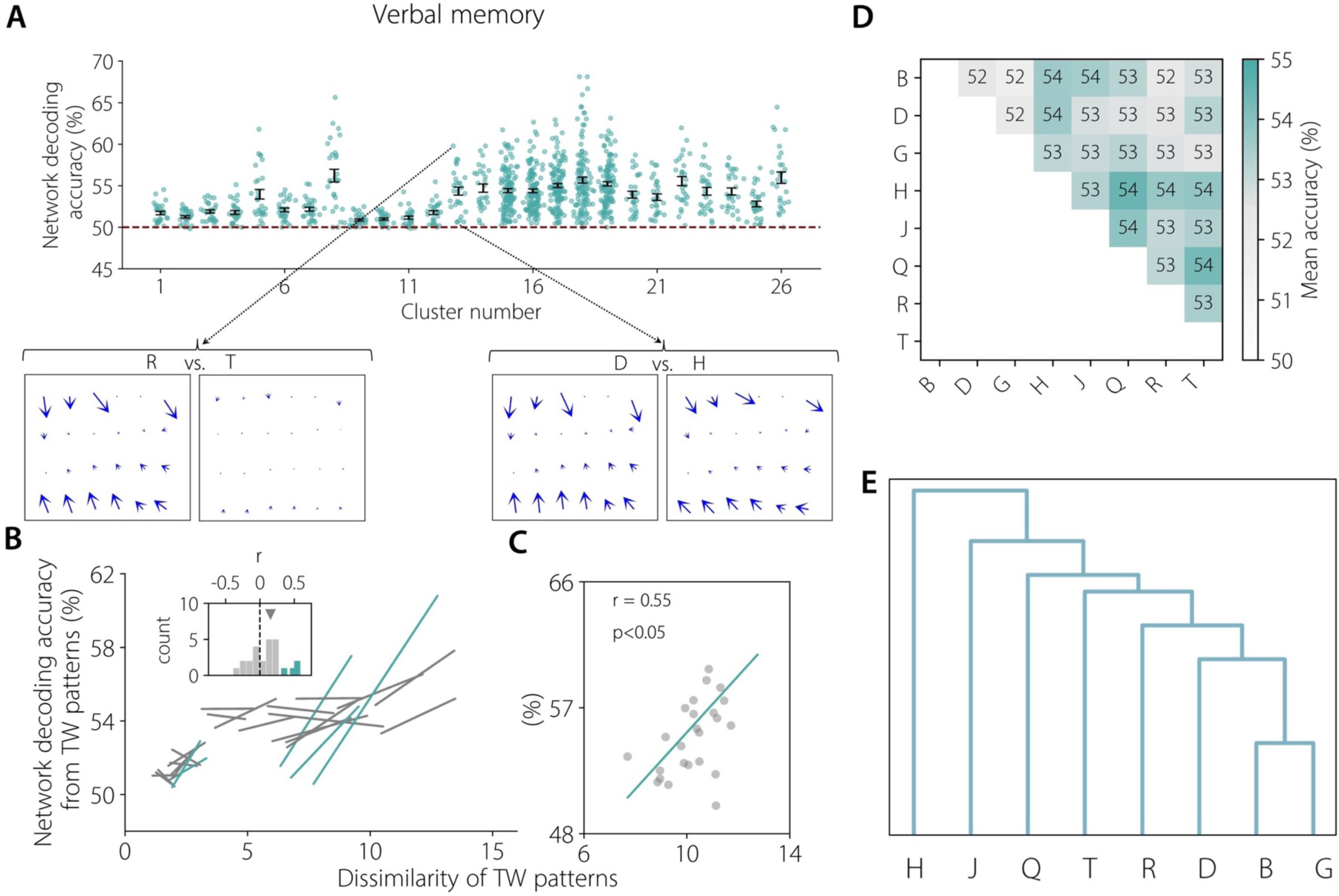
Decoding specific cognitive states of human memory representations from traveling waves. **(A) Network decoding accuracy for each oscillation cluster in the verbal memory task.** Each point in the plot corresponds to a pair of specific cognitive states (for example, letter “B” versus letter “D”). Error bars show SEM decoding accuracies across all pairs. Dotted red line denotes chance level (50%). Inset shows that pairs of letters which had higher decoding accuracy had wave patterns that were visually more distinct from each other. Example waves belong to patient #16, mode #3, CF ∼ 20.0 Hz. Note that clusters 13-26 showed higher decoding accuracies compared to clusters 1-12 because of higher number of electrodes (and hence higher number of modes) in their grids (median number of electrodes = 42 in clusters 13-26 compared to median number of electrodes = 24 in clusters 1-12). **(B, C) Network decoding accuracy versus dissimilarity of traveling wave patterns, in the verbal memory task.** Decoding accuracy was mostly positively correlated with the separability of letters, demonstrating that the decoding results are consistent with the results obtained from the MANOVA analysis. One line was fit for each oscillation cluster and the inset shows the histogram of Pearson correlation values across clusters. Pearson correlation values which were statistically significant (*ps* < 0.05) are denoted in green for both the fitted line and the inset plot. **C** shows fitted line for an example oscillation cluster, with each point in the plot denoting a pair of letters. **(D) Mean decoding accuracy across all oscillation clusters for each pair of letters. (D) Dendrogram of D, demonstrating that letter “H” was the most decodable, followed by the letters “J” and “Q”, compared to the other letters.**

## Discussion

We used direct human brain recordings to investigate directional propagation of traveling waves and probe their link to behavior. We found that diverse patterns of traveling waves including rotational, concentric, and complex wave patterns changed their direction and/or strength to distinguish various cognitive states. These results support the notion that the human brain exhibits complex spatiotemporal patterns of traveling waves that reflect complex cognitive processes including memory and specific brain states. Notably because task-related traveling waves often show complex propagation patterns, it indicates that the brain exhibits new types of functionally relevant spatial organizations, extending beyond currently known anatomical and functional hierarchies (Buckner & Krienen, 2013; De Martino et al., 2018). Further, the diverse propagation patterns of these cognition-related traveling waves is consistent with a rich set of models suggesting a role for spatially organized neural assemblies and oscillations in the computational processes underlying memory, cognition, and other behaviors (Freeman, 2003; Pinotsis et al., 2023; Pinotsis & Miller, 2023; Rubino et al., 2006).

### Roles for rotational and concentric traveling waves for human memory processing

Traveling waves have previously been seen in rodents and non-human primates (Agarwal et al., 2014; Aggarwal et al., 2022; Besserve et al., 2015; Bhattacharya, Donoghue, et al., 2022; Davis et al., 2020; Gabriel & Eckhorn, 2003; Hamid et al., 2021; Lubenov & Siapas, 2009; Muller et al., 2014; Nauhaus et al., 2009; Patel et al., 2012; Rubino et al., 2006; Rule et al., 2018; Stroh et al., 2013; Vinck et al., 2010; Zanos et al., 2015), including complex patterns of traveling waves (Liang et al., 2023; Liang et al., 2021; Townsend & Gong, 2018; Townsend et al., 2015; Townsend et al., 2017). These complex wave patterns in animals are also functionally relevant. Rotational waves play a critical role in the prefrontal cortex of non-human primates during working memory tasks, demonstrating a dynamic modulation of wave orientation that aligns with task intervals (Bhattacharya, Brincat, et al., 2022). Additionally, the systemic administration of anesthetics reorganizes these wave patterns; for example, propofol induces more structured low-frequency waves while disrupting the organization and directionality of higher frequency waves, delineating a selective, frequency-specific alteration of cortical dynamics (Bhattacharya, Donoghue, et al., 2022). In rodents, cortex-wide spiral waves not only exhibit mirrored patterns between hemispheres but also between sensory and motor cortices, reflecting the structural and functional symmetry of long-range axonal projections (Ye et al., 2023). We found that, in addition to planar waves, the human cortex shows rotational and concentric wave patterns as well as complex spatial patterns during memory processing. Crucially, these complex wave patterns were widespread, appearing for both low and high frequency oscillations and across left and right hemispheres in both tasks.

Notably we found that rotational wave patterns were especially pronounced during the spatial episodic memory task and in the beta frequency band. Rotational traveling waves have been previously detected during sleep spindles in the human brain, indicating that this type of waves putatively supports complex cognitive processes such as memory consolidation and synaptic plasticity (Muller et al., 2016). Consistent with our hypothesis, we found a greater prevalence of rotational traveling waves during the more cognitively demanding spatial episodic memory task, which involves richer types of information processing compared to the verbal memory task. This putatively suggests a general role of rotational wave patterns for linking widespread brain regions to support complex cognition.

A notable finding in our work was that in addition to the low frequency theta/alpha traveling waves, there were traveling waves in the beta frequency band during memory processing. In our previous work, using information theoretic analysis, we had found greater bottom-up information flow from the hippocampus to the prefrontal cortex in low frequency delta–theta band and higher top-down information flow from the prefrontal cortex to the hippocampus in the beta band during spatial and verbal memory tasks (Das & Menon, 2021, 2022). The high frequency traveling waves that we found may putatively contribute to top-down information flow for transition and maintenance of latent neuronal ensembles into active representations in the hippocampus as has been hypothesized before (Engel & Fries, 2010; Spitzer & Haegens, 2017) whereas, the low frequency waves may putatively be related to hippocampal signaling of pattern completion associated with memory processing that is conveyed to multiple prefrontal and parietal regions (Eichenbaum, 2017). Traveling waves also exist in the hippocampus during memory processing (Zhang & Jacobs, 2015), thus future studies may wish to identify links between subcortical waves with the cortical waves that we focus on here. An additional future area of work may be to compare the relation between memory-related traveling waves with similar signals at even slower frequencies, as recent fMRI studies have detected waves of hemodynamic activities at slower frequencies during naturalistic language tasks and resting state (Bolt et al., 2022; Raut et al., 2021).

An interesting aspect of our results concerns concentric source and sink waves. Higher prevalence of source waves has been previously detected during sleep spindles in the human brain, indicating that sources may putatively support complex cognitive processes such as memory consolidation and synaptic plasticity compared to sinks (Muller et al., 2016). Because they involve waves propagating outward from a single location, source waves would putatively indicate that a small local group of neuronal assemblies dominate information flow by routing their information outward in a direction where they flow towards widespread brain areas. Consistent with our hypothesis, we found that, whereas in the more complex spatial memory task there are more sources compared to sinks (7 sources versus 2 sinks), in the simpler verbal memory task there are more sinks compared to sources (7 sources versus 15 sinks). Inversely, because the propagating waves converge on a single location, the cortex at the center of sink waves globally integrates information from distributed cortical networks to a specific set of neuronal ensembles for integration and potential memory binding. Going forward, it may be useful to compare the relative prevalence of source versus sink waves in various tasks to reveal variations in the extent of cortical integration (sinks) versus propagation (sources).

### Possible mechanisms for complex patterns of traveling waves

Our findings converge well with theoretical predictions from biologically plausible neural models based on weakly coupled oscillators (Bhattacharya et al., 2021; Sato, 2022). These models hypothesize that complex patterns of waves can be generated locally based on the initial spatial activation of neurons, where each neuron is connected to a few of its neighbors, with distance dependent axonal delays in the order of conduction along unmyelinated horizontal fibers (Davis et al., 2021; Destexhe, 1994; Ermentrout & Kleinfeld, 2001). These locally generated waves can propagate across widespread regions between locally connected neurons and interact with other locally generated waves, to generate complex patterns of propagating oscillations (Bao & Wu, 2003; Huang et al., 2010; Jeong et al., 2002; Schiff et al., 2007). In addition to the coupling functions, a key determinant of the shape of these wave patterns is the presence of local shifts in the amplitude and frequency of local oscillations. In locally coupled oscillator networks, waves tend to propagate away from the cortical locations with the fastest intrinsic oscillation frequencies, following a gradient to the locations with the slowest oscillations (Kopell & Ermentrout, 1990). Thus, local activations of strong or fast oscillations can have a strong influence on the global topography of these traveling waves (Bhattacharya et al., 2021; Huang et al., 2010; Sato, 2022). Using this model, based on the spatial location of the locally generated wave sources and their relative frequencies, a wide range of complex wave patterns such as spirals and spatially heterogeneous wave patterns can be generated (Kopell & Ermentrout, 1990).

A surprising aspect of our findings is the large degree of heterogeneity we found in the directions and strengths of the task-related traveling waves, across oscillation clusters, brain regions, and subjects. This degree of variability is in line with recent advances in structural MRI studies, which have shown that inter-individual variability can have a major impact on human behavior (Genon et al., 2022; Kanai & Rees, 2011). This type of variability has also been shown in functional MRI studies, which found that inter-individual variability in the functional connectivity of task-related activations correlates with the underlying inter-individual variability in anatomical connectivity (Mueller et al., 2013). In the same way that individual differences in structural patterns drive large-scale differences in fMRI signals, an interesting area of future work will be to assess the degree to which broad inter-subject differences in traveling waves may be driven by small anatomical differences across individuals.

It is notable that the traveling waves that we detected often traveled through the sulci and gyri, and even across the Sylvian fissure, across distributed brain areas. For example, for mode 3 in **Figure 6A**, the traveling wave propagated through the Sylvian fissure. Recent work on computational modeling of neural field theory of brain waves (Anastassiou et al., 2011; Davis et al., 2021; Pinotsis & Miller, 2022, 2023) has shown that stable electric fields are capable of carrying information through sulci and gyri, thus providing a mechanism for ephaptic coupling of multiple, distributed brain areas. Moreover, one previous fMRI study on traveling waves has found long-range ephaptic coupling through the sulci of widespread brain areas (Xu et al., 2023). The traveling waves that we observed in our tasks could putatively be a manifestation of these stable electric fields and their interactions with the sulci and gyri, which allows for ephaptic coupling between distributed brain areas (Ermentrout & Kleinfeld, 2001). The focus of the current work was on the analysis of cortical traveling waves, however, recent computational modeling work (Bhattacharya et al., 2021) and experimental data in rodents (Ye et al., 2023) have suggested that traveling waves can exist in the subcortical brain areas such as the thalamus, and can have a major impact on the cortical propagating waves. This also hints at the possibility of three-dimensional traveling waves coordinating neural activity between the cortical and subcortical brain areas, rather than waves propagating only along the cortical surface. Denser sampling of depth electrodes in three-dimension may help to better understand the role of complex patterns of traveling waves in human behavior.

### Behavioral relevance of diverse patterns of traveling waves

Lesion (Bohbot et al., 1998; Maguire et al., 1996; Parto Dezfouli et al., 2021; Spiers et al., 2001) and electrophysiology (Boran et al., 2019; Jacobs et al., 2013; Johnson et al., 2018; Johnson et al., 2017; Miller et al., 2018; Stangl et al., 2021; Stevenson et al., 2018) studies in humans have shown prominent involvement of widespread bihemispheric brain areas spanning the frontal, temporal, and parietal cortices in spatial and verbal memory processing. Consistent with this, our findings indicate that complex spatial patterns of traveling waves are present in both hemispheres at roughly similar levels. Even more intriguingly, we found a critical role for the complex patterns of traveling waves during the distractor period of the spatial memory task. Our results showed that the wave strength of the complex wave is the highest during the distractor period compared to other task periods such as encoding, navigation, retrieval, etc. This suggests more localized information processing during the distractor period compared to other task periods. Future studies with varying load during the distractor period are necessary to probe the role of these complex traveling waves in the human brain.

In many ways, during behavior, cortical information flow follows a hierarchy in which neural activity associated with sensory processes flows “forward” towards other cortical areas, while that related to higher order cognitive processes such as memory retrieval feeds “backward” from frontal cortical areas to coordinate and reinstate neural activity in other brain regions (Friston, 2008; Rabinovich et al., 2012; Vezoli et al., 2021). Our earlier study (Mohan et al., 2024) and others (Alamia & VanRullen, 2019) showed a role for these forward and backward patterns in traveling waves. However, going further, our present results show an additional new hierarchy of traveling waves, by showing more complex spatial patterns which appear to complement this feedforward-feedback cortical hierarchy. Our current work converges on a large body of recent work on traveling waves which has found that complex patterns of waves in rodents and non-human primates (Bhattacharya, Brincat, et al., 2022; Bhattacharya, Donoghue, et al., 2022; Liang et al., 2023; Liang et al., 2021; Townsend et al., 2015) as well as in humans (Muller et al., 2016), play a critical role in cognition. Since traveling waves are known to be closely associated with spiking activity of neurons (Davis et al., 2020), their propagation putatively reflects packets of neuronal activity sequentially scanning distributed brain areas to transiently reorganize functional connectivity between them, to represent complex behaviors in human memory processing (Eichenbaum, 2000; Mesulam, 1990). This would indicate that, whereas the planar waves that we detected in our tasks globally route neural information by directionally propagating to different cortical areas in a feedforward-feedback manner, the rotational waves revisit the same brain areas in multiple cycles, therefore putatively dynamically strengthening functional connectivity between large-scale neuronal assemblies for efficient memory processing, similar to the rotational waves observed during sleep spindles (Muller et al., 2016).

The more complex, heterogenous patterns of waves that we observed putatively route information in more flexible ways by propagating in several directions to rapidly reorganize functional connectivity between neuronal assemblies, to distinguish both broad and specific cognitive states. Recent imaging studies in rodents (Benisty et al., 2024) as well as humans (Demertzi et al., 2019) support this hypothesis by showing that dynamic and complex patterns of functional connectivity can support distinct behaviors independent of the anatomical connectivity. Recent theoretical and computational models have shown that traveling waves play a critical computational role in the visual system by enabling continuous predictions of visual scenes many frames in the future from dynamic and naturalistic visual inputs (Benigno et al., 2023) and may serve general computational roles in complex cognition beyond just visual processing (Pinotsis et al., 2023). Our findings of traveling waves that differentiated cognitive states are thus consistent with this proposal that the complex spatial patterns of waves across the cortex play a role in supporting different computational processes during cognition.

## Conclusions

We used human electrocorticographic recordings and multiple memory tasks to investigate the large-scale electrophysiological basis of distinct cognitive representations subserving human memory processing. We showed that lower frequency theta/alpha and higher frequency beta oscillations are widespread in the neocortex. Using a localized circular–linear regression approach and then independent component analysis, we found that in addition to planar waves, rotational, concentric, and complex patterns of traveling waves are widespread in the human brain. These diverse patterns of traveling waves were able to distinguish both broad and specific cognitive states underlying human memory, and crucially, we were able to robustly decode the cognitive states at the individual subject level by using the features of the propagating traveling waves using machine learning approaches, a significant advance over prior work involving traveling waves. Our methods presented here provide a general approach to the analysis of traveling waves and their association with human behavior and are applicable to other data modalities such as scalp EEG, magnetoencephalographic, optical, as well as fMRI recordings. Fundamentally, these findings are important because they suggest that parallel distributed processing, paced through rhythmic propagating waves with different shapes that flexibly vary according to the current behavior, underlies human memory representations and allows the brain to flexibly adapt to different task demands subserving complex human behaviors and we quote (Mesulam, 1990).

## Materials and Methods

### Human subjects

We examined direct brain recordings from 24 patients with pharmaco-resistant epilepsy who underwent surgery for removal of their seizure onset zones. Patients who performed the Treasure Hunt (TH) spatial episodic memory task (N=9, 4 females, minimum age = 20, maximum age = 57, mean age = 36.6, see below for details) were part of a larger data collection initiative known as the University of Pennsylvania Restoring Active Memory (UPENN-RAM) project. The direct recordings of these patients can be downloaded from a data sharing archive hosted by the UPENN-RAM consortium (URL: http://memory.psych.upenn.edu/RAM), shared by Kahana and colleagues (Jacobs et al., 2016). Prior to data collection, research protocols and ethical guidelines were approved by the Institutional Review Board at the participating hospitals and informed consent was obtained from the participants and guardians (Jacobs et al., 2016). Recordings from the patients who performed the Sternberg (ST) verbal working memory task (N=15, 7 females, minimum age = 20, maximum age = 58, mean age = 35.9, see below for details), were collected at four hospitals (Thomas Jefferson University Hospital, Philadelphia; University of Pennsylvania Hospital Philadelphia; Children’s Hospital of Philadelphia, and University Hospital Freiburg). The direct recordings of these patients can be downloaded from https://memory.psych.upenn.edu/Data, shared by Kahana and colleagues (Jacobs & Kahana, 2009). Similar to the Treasure Hunt spatial episodic memory task, all patients who performed the Sternberg verbal working memory task consented to having their brain recordings used for research purposes and the research was approved by relevant Institutional Review Boards.

### Electrophysiological recordings and preprocessing

Patients were implanted with different configuration of electrodes based on their clinical needs, which included both electrocorticographic (ECoG) surface grid and strips as well as depth electrodes. In this work, we only examined the ECoG grid electrodes on the cortical surface. ECoG recordings were obtained using subdural grids (contacts placed 10 mm apart) using recording systems at each clinical site. Recording systems included DeltaMed XlTek (Natus), Grass Telefactor, and Nihon-Kohden EEG systems.

Anatomical localization of electrode placement was accomplished by co-registering the postoperative computed CTs with the postoperative MRIs using FSL (FMRIB (Functional MRI of the Brain) Software Library), BET (Brain Extraction Tool), and FLIRT (FMRIB Linear Image Registration Tool) software packages. Preoperative MRIs were used when postoperative MRIs were not available. From these images, we identified the location of each recording contact on the CT images and computed the electrode location in standardized Talairach coordinates.

Original sampling rates of ECoG signals in the TH task were 500 Hz, 1000 Hz, and 1600 Hz. ECoG signals in the TH task were downsampled to 500 Hz, if the original sampling rate was higher, for all subsequent analysis. Original sampling rates of ECoG signals in the ST task were 400 Hz, 512 Hz, and 1000 Hz. Therefore, ECoG signals in the ST task were downsampled to 400 Hz, if the original sampling rate was higher, for all subsequent analysis. We used common average referencing (ECoG electrodes re-referenced to the average signal of all electrodes in the grid), similar to our previous studies on traveling waves (Das et al., 2022; Zhang et al., 2018). Line noise (60 Hz) and its harmonics were removed from the ECoG signals. For filtering, we used a fourth order two-way zero phase lag Butterworth filter throughout the analysis.

### Cognitive tasks

#### (a) Treasure Hunt spatial episodic memory task

The patients performed multiple trials of a spatial memory task in a virtual reality environment developed in Unity3D (Miller et al., 2018; Tsitsiklis et al., 2020), where they played a Treasure Hunt task on a laptop computer at the bedside and controlled their translational and rotational movements through the virtual environment with a handheld joystick. In each task trial, subjects explored a rectangular arena on a virtual 3D beach to reach treasure chests that revealed hidden objects, with the goal of encoding the location of each item encountered (**Figure 1A**). The virtual beach (100 × 70 virtual units, 1.42: 1 aspect ratio) was bounded by a wooden fence on each side. The locations of the objects changed over the trials, but the shape, size and appearance of the environment remained constant throughout the sessions. The task environment was constructed so that the subject would perceive one virtual unit as corresponding to approximately 1 foot in the real world. Subjects viewed the environment from the perspective of cycling through the environment and the elevation of their perspective was 5.6 virtual units. Each end of the environment had unique visual cues to help the subjects navigate.

Each trial of the task begins with the subject being placed on the ground at a randomly selected end of the environment. The subject initiates the trial with a button press, then navigates to a chest using a joystick. Upon arrival at the chest, the chest opens and either reveals an object, which the subject should try to remember, or is empty. The subject remains facing the open chest for 1.5 sec (*encoding period*) and then the object and chest disappear, which indicates that the subject should navigate (*navigation period*) to the next chest that has now appeared in the arena. Each trial consists of four chests; two or three (randomly selected, so that subjects could not predict whether the current target chest contained an object, which served to remove effects of expectation and to encourage subjects to always attend to their current location as they approached a chest) of the chests contain an object, and the others are empty. Each session consists of 40 trials, and in each session, subjects visit a total of 100 full chests and 60 empty chests. The chests are located pseudorandomly throughout the interior of the environment, subject to the restrictions that no chest can be placed within 11 virtual units of another and that all chests must be at least 13 virtual units from the arena boundary. This 11 virtual unit restriction ensures that chest locations are varied in a trial. There are no constraints based on previous trials, and all object identities are trial-unique and never repeated within a session. After reaching all four chests of a trial, subjects are transported automatically so that they have an elevated 3/4 overhead perspective view of the environment at a randomly selected end of the environment. The reason for this perspective shift was to speed up the recall period, preserving patient testing time to provide a relatively larger number of memory encoding events. They then perform a distractor task (*distractor period*), a computerized version of the “shell game”, before entering the *retrieval period*. During recall, subjects are cued with each of the objects from the trial in a random sequence and asked to recall the location of the object. In each recall period, they first indicate their confidence (*confidence period*) to remember the location of the object (“high”, “medium”, or “low”). Next, they indicate the precise location of the object by placing a cross-hair at the location in the environment that corresponds to the location of the cued item. After the location of each object of the trial is indicated, the feedback stage (*feedback period*) of each trial begins. Here, subjects are shown their response for each cued object in the trial, via a green circle if the location was correct and a red circle if it was incorrect. Subjects receive feedback on their performance, following a point system where they receive greater rewards for accurate responses. A response is considered correct if it is within 13 virtual units of the true object location. Mean accuracy across subjects was ∼41%.

We analyzed the 1.5 sec long trials from the encoding periods of the TH task. For the navigation periods, we analyzed 1.5 sec long time segments approximately corresponding to the middle of the navigation trial. Similarly, for the distractor periods, we analyzed 1.5 sec long time segments approximately corresponding to the middle of the distractor trial. For the confidence periods, we analyzed 1.5 sec recording immediately following the presentation of the visual cues. For the retrieval periods, we analyzed 1.5 sec recording immediately prior to the retrieval of the objects. For the feedback periods, we analyzed 1.5 sec time segments immediately following the feedback.

#### (b) Sternberg verbal working memory task

Patients performed multiple trials of a Sternberg verbal working memory task (Jacobs & Kahana, 2009; Sternberg, 1966). In each trial of the task, patients were presented with a list of one to three English letters on the screen of a bedside laptop computer (**Figure 1B**). During this presentation portion of the trial, first a fixation cross appeared, and then the letters were displayed sequentially on the computer screen. Each letter appeared on screen for 1 sec. Patients were instructed to closely attend to each stimulus presentation and to silently hold the identity of each item in memory. The letter lists included only consonants (i.e., no vowels) to prevent patients from using mnemonic strategies (e.g., remembering the entire list as an easily pronounceable word-like sound). After the presentation of each list, the response period began when a probe item was displayed. Then patients responded by pressing a key to indicate whether the probe was present in the just-presented list or whether it was absent. After the key press, the computer indicated whether the response was correct, and then a new list was presented. Across all letter presentations, patients viewed an 8 or 16 letter subset. Mean accuracy across subjects was ∼92%. We analyzed the 1 sec long trials from the encoding period of each letter.

### Identification of oscillations

To characterize propagating traveling waves in an ECoG grid, we first identified narrowband oscillations at theta/alpha and beta bands. We adopted methods similar to our previous approach (Das et al., 2022). An advantage of this algorithm is that it accounts for several complexities of human brain oscillations measured with ECoG signals, including differences in electrode positions across subjects and variations in oscillation frequencies across individuals. Identifying oscillation frequency is crucial since, by definition, a traveling wave involves a single frequency and whose phase progressively propagates through the ECoG electrodes, thus making it possible to detect the traveling wave when it passes by these electrodes.

We first used Morlet wavelets to compute the power of the neural oscillations at 200 frequencies logarithmically spaced from 3 to 40 Hz. To identify narrowband oscillations at each electrode, we fit a line to each patient’s mean power spectrum (mean across trials) in log–log coordinates using robust linear regression (Das et al., 2022) (**Figure 1D**). We then subtracted the actual power spectrum from the regression line. This normalized power spectrum removes the 1/f background signal and emphasizes narrowband oscillations as positive deflections (**Figure 1E**). We identified narrowband peaks in the normalized power spectrum as any local maximum greater than one standard deviation above the mean (**Figure 1E**). We repeated this procedure for each of encoding, confidence, navigation, retrieval, distractor, and feedback periods in the TH task and also for the English letters in the ST task. The mean of the peak frequencies of the electrodes of a ECoG grid was defined as the cluster frequency (CF). Since by definition, a traveling wave involves an oscillation frequency, we only included oscillation clusters for which at least 2/3^rd^ of the electrodes in the ECoG grid had a narrowband peak in the power spectrum, for further analyses, to ensure that the traveling waves are mostly driven by oscillatory activity. We excluded 5 patients in the TH task and 2 patients in the ST task from further analyses since these patients did not meet this criterion. This resulted in 9 patients in the TH task and 15 patients in the ST task to be included in all subsequent analyses below. Overall, ∼87% of ECoG electrodes in the TH task and ∼86% of ECoG electrodes in the ST task had a narrowband oscillation in one of the theta/alpha or beta frequency bands, indicating that an overwhelming number of ECoG electrodes show oscillatory activity. Note that in many cases, electrodes in the same ECoG grid showed multiple peaks in the power spectrum, i.e, they had peaks at both the theta/alpha and beta frequency bands. We estimated traveling waves for each of these narrowband frequencies separately.

### Identification of traveling waves

We next estimated traveling waves corresponding to each of the oscillation clusters identified above. A traveling wave can be described as an oscillation that moves progressively across a region of cortex. Quantitatively, a traveling phase wave can be defined as a set of simultaneously recorded neuronal oscillations at very similar frequencies whose instantaneous phases vary systematically with the locations of the recording electrodes. We used a localized circular-linear regression approach, assuming that the relative phases of the oscillation clusters exhibit a linear relationship with electrode locations *locally* (Das et al., 2022). This locally circular-linear fitting of phase-location can detect complex patterns (Ermentrout & Kleinfeld, 2001; Muller et al., 2016) of traveling waves in an oscillation cluster in addition to planar traveling waves.

To identify traveling waves from the phases of each oscillation cluster, we first measured the instantaneous phases of the signals from each electrode of a given cluster by applying a 4^th^ order Butterworth filter at the cluster’s oscillation frequency (bandwidth [f_p_ ×.85, f_p_ / .85] where f_p_ is the peak frequency). We used Hilbert transform on each electrode’s filtered signal to extract the instantaneous phase.

We used circular statistics to identify traveling waves for each oscillation cluster at each time point (Fisher, 1993). We first projected the 3-D Talairach coordinates for each cluster into the best-fitting 2-D plane using principal component analysis (PCA). We projected the electrode coordinates into a 2-D space to simplify visualizing and interpreting the data. For each spatial phase distribution, we then used two-dimensional (2-D) localized circular–linear regression to assess whether the observed phase pattern varied linearly with the electrodes’ coordinates in 2-D. In this regression, for each electrode in a given oscillation cluster, we first identified the neighboring electrodes that were located within 25 mm distance of the given electrode, constituting a sub-cluster of the given cluster. Let *x_i_* and *y_i_* represent the 2-D coordinates and *θ_i_* the instantaneous phase of the *i*th electrode in a sub-cluster.

We used a 2-D circular-linear model

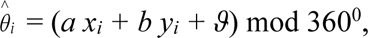

where 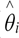 is the predicted phase, *a* and *b* are the phase slopes corresponding to the rate of phase change (or spatial frequencies) in each dimension, and *ϑ* is the phase offset. We converted this model to polar coordinates to simplify fitting. We define *α* = atan2(*b, a*) which denotes the angle of wave propagation and 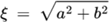 which denotes the spatial frequency. Circular–linear models do not have an analytical solution and must be fitted iteratively (Fisher, 1993). We fitted *α* and ξ to the distribution of oscillation phases at each time point by conducting a grid search over *α* ∈ [0^0^, 360^0^] and ξ ∈ [0, 18]. Note that ξ = 18 corresponds to the spatial Nyquist frequency of 18^0^/mm corresponding to the spacing between neighboring electrodes of 10 mm.

In order to keep the computational complexity tractable, we used a multi-resolution grid search approach. We first carried out a grid search in increments of 5^0^ and 1^0^/mm for *α* and *ξ*, respectively. The model parameters (*a= ξ*cos(*α*) and *b= ξ*sin(*α*)) for each time point are fitted to most closely match the phase observed at each electrode in the sub-cluster. After having relatively coarse estimates of *α* and *ξ*, we then carried out another grid search in increments of 0.05^0^ and 0.05^0^/mm around a ± 2.5^0^ and ± 0.5^0^/mm neighborhood of the coarse estimates of *α* and *ξ*, respectively, to have refined estimates of *α* and *ξ*. We computed the goodness of fit as the mean vector length *r̅* of the residuals between the predicted (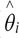) and actual (*θ_i_*) phases (Fisher, 1993),

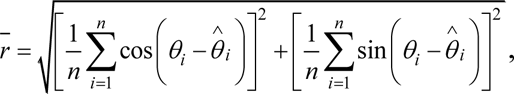

where *n* is the number of electrodes in the sub-cluster. The selected values of *α* and ξ are chosen to maximize *r̅*. This procedure is repeated for each sub-cluster of a given oscillation cluster. To measure the statistical reliability of each fitted traveling wave, we examined the phase variance that was explained by the best fitting model. To do this, we computed the circular correlation between the predicted (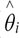) and actual (*θ_i_*) phases at each electrode:

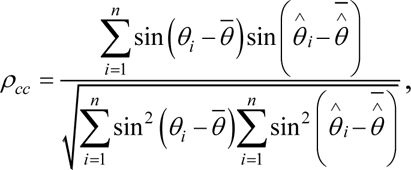

where bar denotes averaging across electrodes. We refer to 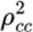 as the wave strength (Das et al., 2022) as it quantifies the strength of the traveling wave (note that 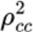 has been referred to as the phase gradient directionality (PGD) in some prior studies (Muller et al., 2016; Rubino et al., 2006; Zhang et al., 2018)). Other features of traveling waves such as the wavelength (2π/spatial frequency) and the speed (wavelength × frequency) can be readily derived from the parameters of the above 2-D model. Note that traveling waves in some prior studies were detected and analyzed by calculating the spatial gradient of the phases of the recordings from ECoG electrodes (Halgren et al., 2019; Muller et al., 2016), however, phase gradients can only be calculated in two directions (forward and backward), so only a subset of neighboring electrodes of a given electrode are included in these analyses of spatial gradient. Since our approach directly includes all possible neighboring electrodes (termed as a sub-cluster in our analysis) in the circular-linear regression model, it results in a more efficient estimate of the traveling waves parameters.

We note that since a few of the ECoG electrodes did not have a narrowband oscillation, we estimated the traveling waves for those electrodes by an extrapolation procedure where, for the given electrode, we substituted the mean of the traveling waves of all electrodes within a 25 mm radius of the electrode under consideration. This extrapolation step was necessary for classification of each oscillation cluster as one of the wave categories using curl and divergence analysis (rotational or expanding/contracting or complex, see section below on ***Identification of rotational and concentric traveling waves***). This is reasonable since as mentioned above, an overwhelming number of ECoG electrodes showed oscillatory activity (∼87% of ECoG electrodes in the TH task and ∼86% of ECoG electrodes in the ST task had a narrowband oscillation in one of the theta/alpha or beta frequency bands). Nevertheless, we reran our ICA analysis (see section below on ***Identification of modes using complex independent component analysis (CICA)***) and statistical significance using MANOVA (see section below on ***Statistical analysis***) without the extrapolation step, and found that the same modes and clusters were still statistically significant in both the TH and ST tasks, indicating that the extrapolation step has very minimal effect on the results reported here.

### Identification of stable epochs

Since a traveling wave is composed of phase patterns that vary relatively smoothly across space and time, we sought to characterize the spatiotemporal stability of the traveling waves that we detected in our localized circular-linear regression approach above. We defined *stability* as the negative of the mean (across all electrodes in an ECoG grid) of the absolute values of the difference between the direction and strength of traveling waves observed at consecutive time-points, defined as

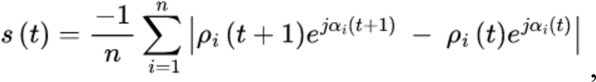

where, *ρ_i_* denotes the strength of the traveling wave for the *i*th electrode, *α_i_* denotes the direction of the traveling wave for the *i*th electrode, *n* denotes the number of electrodes in an ECoG grid, *s* denotes stability, *t* denotes time, and denotes square root of minus one. We repeated this procedure for each pair of consecutive time-points and for each trial of the encoding, retrieval, navigation, etc. periods in the TH task and each trial of the English letters in the ST task. Stability for each trial was z-scored. We identified stable epochs as those for which all stability values were above a predefined threshold. We chose this threshold to be the mean (which is zero since the stability values are z-scored) of the stability values for a given trial. We ran our stability analysis across all trials and detected stable epochs corresponding to each of the encoding, retrieval, navigation, etc. periods in the TH task and each of the English letters in the ST task (**Figures 1F, G**).

Previous research in rabbit field potential recordings (Freeman & Rogers, 2002; Freeman & Schneider, 1982) have found that theta frequency traveling waves last ∼80-100 msec in duration. Moreover, large-scale, whole-brain computational modeling in humans using neural field theory have shown that spatiotemporally stable traveling waves last ∼50-60 msec in duration (Roberts et al., 2019). Therefore, we rejected all the stable epochs which were less than 25 time-points long, from further analysis; these short-length stable epochs are putatively related to noise rather than cognition (Roberts et al., 2019). This corresponded to 50 msec in the TH task (*fs* = 500 Hz) and 62.5 msec in the ST task (*fs* = 400 Hz).

We also calculated the histograms of the occurrence (estimated by aggregating all time-points at which these stable epochs appeared) of these stable epochs, however, we did not find any strong temporal modulation of these stable epochs during the task periods (**Supplementary Figure 3**).

### Identification of modes using complex independent component analysis (CICA)

Since the direction and strength of the traveling wave remains almost the same at each time-point of a stable epoch identified above, we averaged the direction and strength of the traveling waves across all time points for each electrode and each stable epoch. In this way, we find one wave pattern associated with each stable epoch. We concatenated the wave patterns (i.e., direction and strength) for all stable epochs across encoding, retrieval, navigation, etc. periods into a single matrix and then passed this matrix as input to the complex version of the independent component analysis (CICA) (Fu et al., 2015; Li & Adalı, 2010). The complex version of the ICA was necessary, as compared to the real version of the ICA, to incorporate the 2-D directions of the traveling waves, weighted by the strength (*ρ*cos(*α*) and *ρ*sin(*α*)), defined for each electrode in the ECoG grid. We then extracted the independent activation functions (or, weights) (each activation function corresponds to one of the stable epochs) and the corresponding modes (“raw modes” in **Figure 2**) as the output from the ICA (Fu et al., 2015; Li & Adalı, 2010). Multiplication of each of these modes with the mean of the weights across all stable epochs corresponding to that specific mode results in a unique wave pattern (“mean modes” in **Figure 2**) associated with that mode. At the individual epoch level, a higher ICA weight for that epoch corresponding to a specific mode indicates higher representation of that wave pattern in that specific epoch and a lower ICA weight for an epoch corresponding to a specific mode indicates lower representation of that wave pattern in that specific epoch (**Figure 2**). Moreover, the higher the variance explained by a given mode, the higher will be its representation across the trials (also known as a scree plot, see **Figure 4E**). In this way, we can extract the ICA weights for each of the encoding, retrieval, navigation, etc. periods in the TH task and each of the English letters in the ST task and directly compare them using statistical significance (for example, encoding vs. retrieval, letter “B” vs. letter “G”, etc.); see ***Statistical analysis*** section for details.

### Identification of planar traveling waves

After extracting the mean modes from the CICA procedure above, we next sought to classify each of the mean modes as one of “planar”, “rotational”, “concentric” (“expanding” or “contracting”), or “complex” categories (**Figure 3, Supplementary Figures 7-16**) (Townsend et al., 2015), to identify global patterns of traveling waves. For detecting planar traveling waves, we calculated the mean wave direction (weighted by the strength of the wave) of all electrodes in an oscillation cluster and compared it with a predefined threshold. For planar wave detection, this threshold was chosen to be 0.6. This threshold was first manually optimized by hit-and-trial method for the TH task (we estimated this threshold to be 0.6) and hence was independent of the ST task. The same threshold was then used for the ST task to detect planar traveling waves. This procedure ensured a relatively unbiased selection of the threshold. Visually, this procedure of threshold selection yielded reasonably good results (**Figure 3**) and small changes in this threshold did not substantially change the results reported here.

### Identification of rotational and concentric traveling waves

In addition to detecting planar traveling waves, we also detected rotational and concentric traveling waves in the mean modes using the *curl* and *divergence* metrics respectively (Muller et al., 2016). Curl can detect rotational patterns (for example, clockwise or counter-clockwise rotation) in wave dynamics and divergence can detect expanding/contracting patterns (for example, source or sink) in wave dynamics (Muller et al., 2016). We first calculated the curl and divergence metrics for each electrode and then calculated the mean curl and divergence across all electrodes in an oscillation cluster. If the mean curl or divergence metrics cross some predefined threshold, then we declare those wave patterns to be rotational or concentric, respectively. This threshold was chosen to be 0.75 for rotational wave detection and 0.4 for concentric wave detection. Similar to the threshold selection for the detection of planar waves, threshold selection for the rotational and concentric waves was first manually optimized by hit-and-trial method for the TH task (we estimated this threshold to be 0.75 for rotational waves and 0.4 for concentric waves, respectively) and hence was independent of the ST task. The same threshold was then used for the ST task to detect rotational and concentric traveling waves. This procedure again ensured a relatively unbiased selection of the threshold. Visually, this procedure of threshold selection yielded reasonably good results for the rotational and concentric waves as well (**Figure 3**) and small changes in this threshold did not substantially change the results reported here.

Waves that could not be classified as one of the planar, rotational, or concentric, were designated as complex waves. Even though there was no global pattern associated with these complex traveling waves, many of these complex waves showed interesting local patterns, where a subset of electrodes in an ECoG grid showed planar, rotational, or concentric waves (**Figure 9, Supplementary Figures 7-16**).

### Decoding analysis

Our final goal was to test whether we could robustly decode the broad and specific cognitive states from the diverse traveling wave patterns in our datasets. Decoding the different cognitive states in our datasets is an example of a multiclass classification problem. We converted this problem into a series of binary classification tasks, as these can be solved straightforwardly with various multivariate algorithms. We trained multilayer neural networks (Bernardi et al., 2020), with cross-validation, for classifying pairwise cognitive states (for example, encoding versus retrieval, letter “D” versus letter “J”, etc.), separately for the spatial and verbal memory tasks. We used a PyTorch-based four-layer neural network for this binary classification problem (Paszke et al., 2019). We used the extracted weights from the ICA procedure as features for training our neural network classifiers. We observed that increasing the number of modes for training increased the network test decoding accuracy (**Figure 11B**). Therefore, we included all the modes for classifying the cognitive states and trained a neural network classifier for each oscillation cluster separately.

The multilayer network architecture comprised an input layer, two hidden layers (32 and 16 neurons respectively), and an output layer with a sigmoid activation function, similar to the neural network architectures previously used for classifying cognitive states (Bernardi et al., 2020). ReLU activation functions were applied to the hidden layers to introduce nonlinearity and improve generalization (Dahl et al., 2013). The network architectures were kept the same across the spatial and verbal memory datasets, and individual models were trained for each oscillation cluster separately to optimize neural network hyperparameters. To further enhance generalization, dropout regularization was implemented after each hidden layer (Srivastava et al., 2014). For model optimization, we used Binary Cross Entropy as the loss function, ideal for binary classification tasks (Ruby & Yendapalli, 2020), and the Adam optimizer from PyTorch during training (Paszke et al., 2019).

To ensure robust optimization and generalization, we employed cross-validation procedures for iterative training over multiple epochs. For each pair of cognitive state, we first rebalanced the data such that we have an equal number of epochs per cognitive state (van Gerven et al., 2013). We used a five-fold cross-validation technique in which the network model was fitted on 80% of the data and then its performance was tested on the remaining 20%. We used a subsampling procedure where each epoch was randomly assigned to a particular fold, subject to the constraint that all cognitive states are evenly represented (van Gerven et al., 2013), and then averaged the network decoding accuracies across folds.

### Statistical analysis

We directly compared the weights estimated from the CICA procedure above between encoding, retrieval, navigation, etc. periods in the TH task and between the English letters in the ST task, for each mode. Since the weights are complex, we used multivariate analysis of variance (MANOVA) to statistically distinguish weights corresponding to different cognitive states using the following model: *Real + Imag ∼ States*, where *Real* and *Imag* are the real and imaginary parts of the weights respectively and *States* are encoding, retrieval, navigation, etc. in the TH task or letters “B”, “G”, etc. in the ST task. We used this model for each mode and applied FDR-corrections for multiple comparisons (*p* < 0.05) across all modes for each oscillation cluster. A statistical significance would indicate that traveling waves shift their direction and/or strength to form distinct directional patterns which can distinguish different behavioral states such as encoding, retrieval, navigation, etc. or letters such as “B”, “G”, etc. We designated an oscillation cluster to be significant if at least one of the modes from the ICA for that cluster showed statistical significance in MANOVA.

We conducted surrogate analysis to test the significance of the estimated stable epochs (see ***Identification of stable epochs*** section above) and whether the observed stable epochs are beyond chance levels. We shuffled the trial labels (encoding, retrieval, etc. in the TH task and “B”, “G”, etc. in the ST task) and electrodes, so that the spatial topography for the corresponding cognitive state is destroyed, and then ran the stable epoch analysis using identical methodology as above. In this way, we built a surrogate distribution by aggregating all time-points corresponding to these shuffled stable epochs against which we then compared the aggregated time-points from the empirical stable epochs (*p* < 0.05).

For assessing statistical significance for the group level results related to fraction of wave strength and fraction of significance (**Figures 4, 7-10**), we used chi-squared tests with FDR-corrections for multiple comparisons (*p* < 0.05) across frequencies (theta/alpha and beta) and hemispheres (left and right).

To estimate the number of significant modes in the CICA output for each cluster, we compared the variance explained for each mode with the theoretical variance threshold 100/*n*, where *n* is the number of electrodes in an ECoG grid. This theoretical variance corresponds to the variance of each mode if the total variance (100%) is equally distributed among all modes. Because of the spatial structure of traveling waves, some modes will explain more variance compared to the other modes in the empirical data (**Figure 4E**). We additionally shuffled the electrodes in each cluster and recalculated the variance distribution across modes and confirmed that the shuffled variance for all modes converged to the theoretical variance threshold of 100/*n*.

Finally, to access group level statistical significance for the decoding accuracy results, we used one-sided sign tests versus chance (*p* < 0.05).

## Acknowledgements

We thank Drs. Uma Mohan and Honghui Zhang for help in traveling waves analysis. This research was supported by an NSF CAREER award to J.J.

## Supplementary File for

### Supplementary tables

**Supplementary Table 1:**
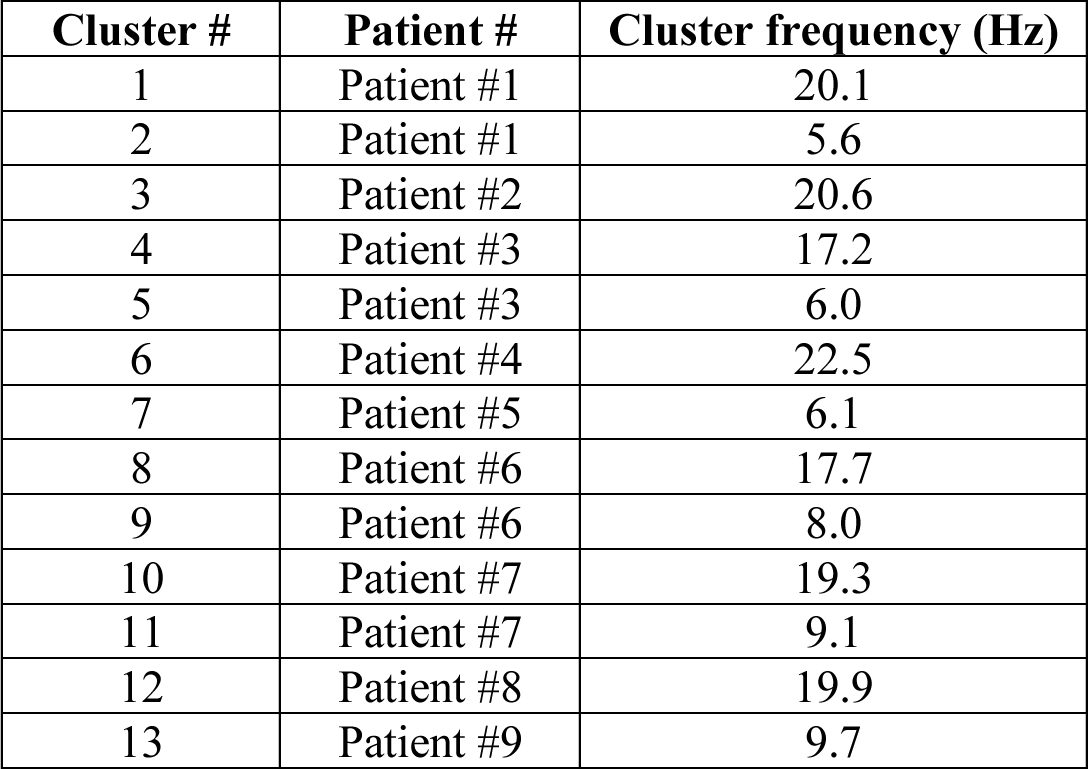
Oscillation clusters in the spatial episodic memory task.

| Cluster # | Patient # | Cluster frequency (Hz) |
| --- | --- | --- |
| 1 | Patient #1 | 20.1 |
| 2 | Patient #1 | 5.6 |
| 3 | Patient #2 | 20.6 |
| 4 | Patient #3 | 17.2 |
| 5 | Patient #3 | 6.0 |
| 6 | Patient #4 | 22.5 |
| 7 | Patient #5 | 6.1 |
| 8 | Patient #6 | 17.7 |
| 9 | Patient #6 | 8.0 |
| 10 | Patient #7 | 19.3 |
| 11 | Patient #7 | 9.1 |
| 12 | Patient #8 | 19.9 |
| 13 | Patient #9 | 9.7 |

**Supplementary Table 2:** Oscillation clusters in the verbal working memory task.

| Cluster # | Patient # | Cluster frequency (Hz) |
| --- | --- | --- |
| 1 | Patient #10, grid #1 | 17.5 |
| 2 | Patient #10, grid #2 | 18.0 |
| 3 | Patient #10, grid #1 | 6.7 |
| 4 | Patient #10, grid #2 | 8.6 |
| 5 | Patient #11 | 19.8 |
| 6 | Patient #12 | 19.9 |
| 7 | Patient #12 | 10.6 |
| 8 | Patient #13 | 6.3 |
| 9 | Patient #14, grid #1 | 25.8 |
| 10 | Patient #14, grid #1 | 8.8 |
| 11 | Patient #14, grid #2 | 9.0 |
| 12 | Patient #15 | 9.1 |
| 13 | Patient #16 | 20.0 |
| 14 | Patient #16 | 9.2 |
| 15 | Patient #17 | 16.0 |
| 16 | Patient #18, grid #1 | 17.3 |
| 17 | Patient #18, grid #2 | 18.1 |
| 18 | Patient #18, grid #1 | 8.7 |
| 19 | Patient #18, grid #2 | 8.3 |
| 20 | Patient #19 | 6.5 |
| 21 | Patient #20 | 5.7 |
| 22 | Patient #21 | 25.4 |
| 23 | Patient #21 | 12.4 |
| 24 | Patient #22 | 11.5 |
| 25 | Patient #23 | 7.0 |
| 26 | Patient #24 | 22.7 |

**Supplementary Table 3:**
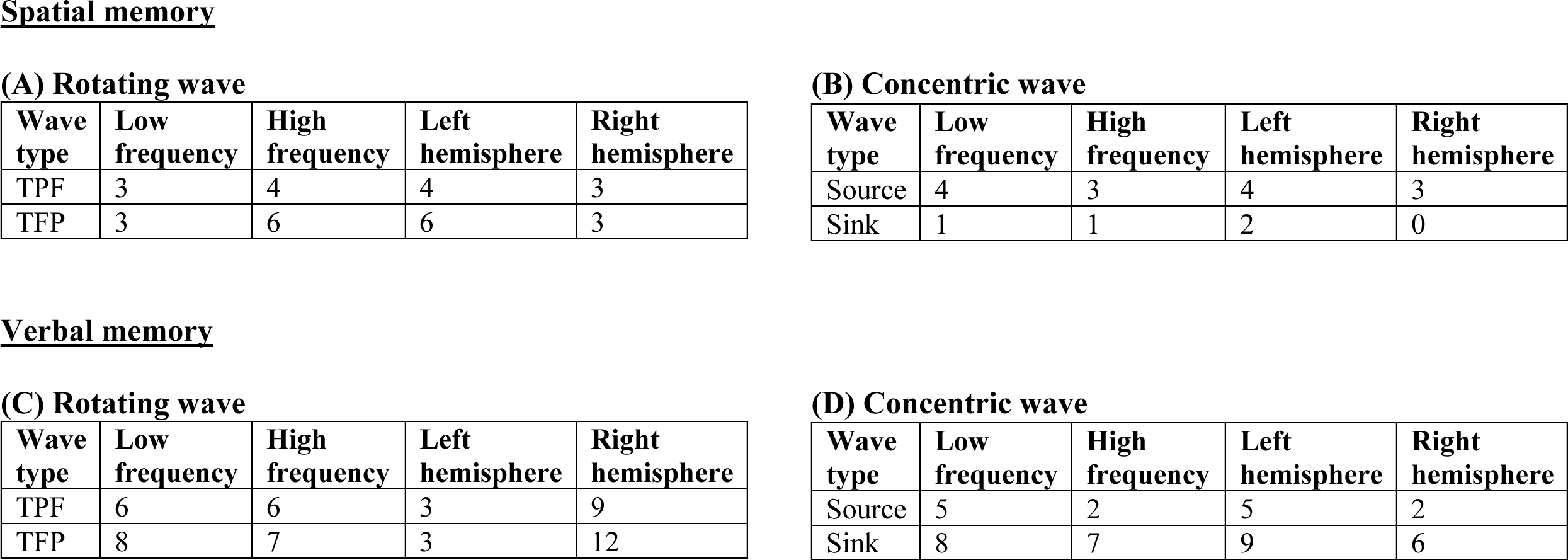
Distribution of each type of rotational (TPF/TFP) and concentric (source/sink) waves for spatial and verbal memory tasks across modes, shown for each frequency and hemisphere. TPF: Temporal-parietal-frontal, TFP: Temporal-frontal-parietal.

**(A) Rotating wave**
| Wave type | Low frequency | High frequency | Left hemisphere | Right hemisphere |
| --- | --- | --- | --- | --- |
| TPF | 3 | 4 | 4 | 3 |
| TFP | 3 | 6 | 6 | 3 |

**(B) Concentric wave**
| Wave type | Low frequency | High frequency | Left hemisphere | Right hemisphere |
| --- | --- | --- | --- | --- |
| Source | 4 | 3 | 4 | 3 |
| Sink | 1 | 1 | 2 | 0 |

**Verbal memory** **(C) Rotating wave**
| Wave type | Low frequency | High frequency | Left hemisphere | Right hemisphere |
| --- | --- | --- | --- | --- |
| TPF | 6 | 6 | 3 | 9 |
| TFP | 8 | 7 | 3 | 12 |

| Wave type | Low frequency | High frequency | Left hemisphere | Right hemisphere |
| --- | --- | --- | --- | --- |
| Source | 5 | 2 | 5 | 2 |
| Sink | 8 | 7 | 9 | 6 |

### Supplementary videos

**Supplementary Video 1:**
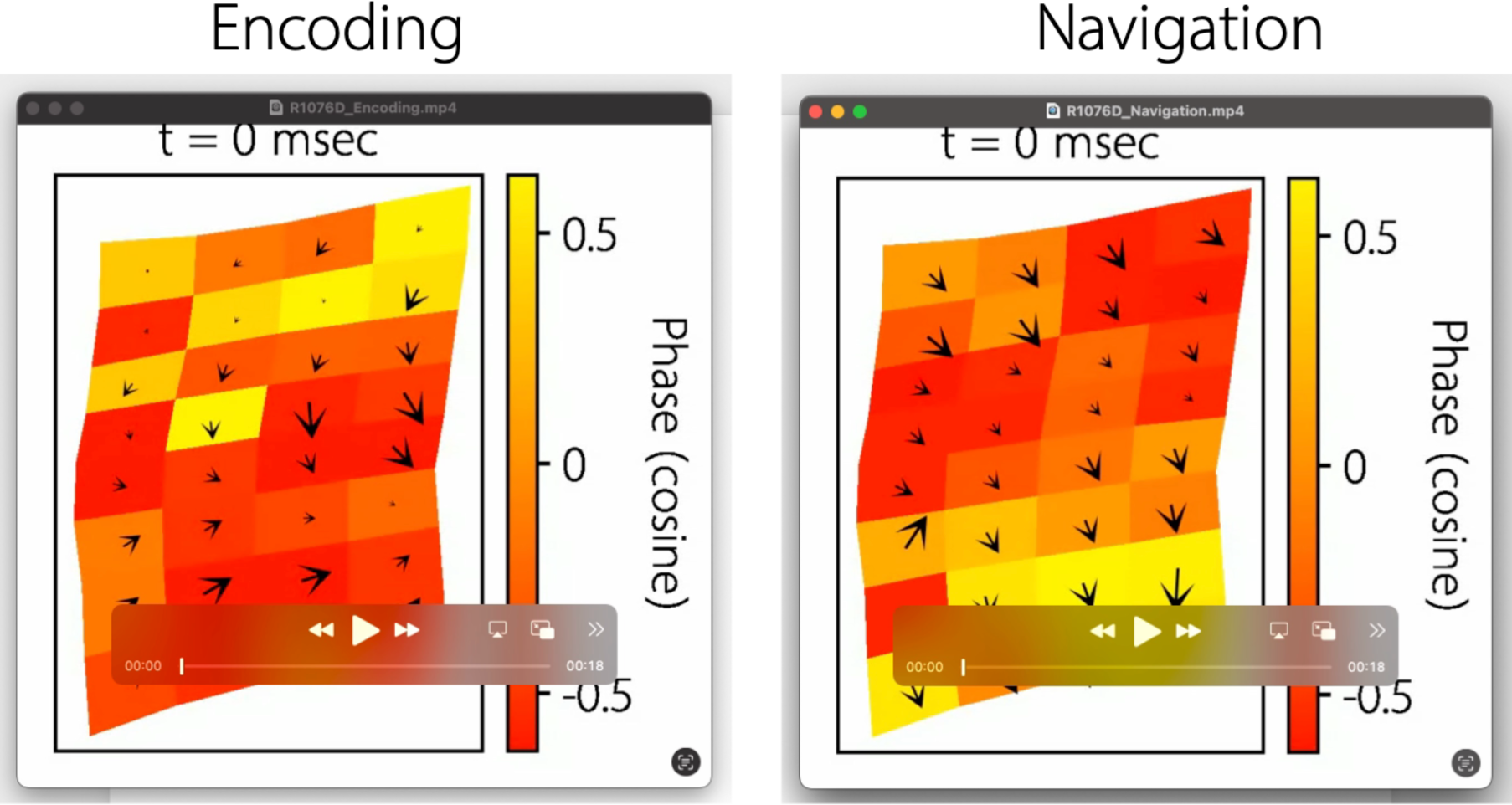
Propagation of traveling waves for two example epochs, one for the Encoding period, and another for the Navigation period, from patient #1, CF ∼ 5.6 Hz. Times denoted refer to the time elapsed since the start of the stable epoch. Note the change in direction of the traveling wave on the lower half of the grid for the encoding versus the navigation period. Shown above are snapshots of the videos. Video link: https://github.com/anupdas777/complex_traveling_waves/blob/main/R1076D_Encoding.mp4 https://github.com/anupdas777/complex_traveling_waves/blob/main/R1076D_Navigation.mp4

**Supplementary Video 2:**
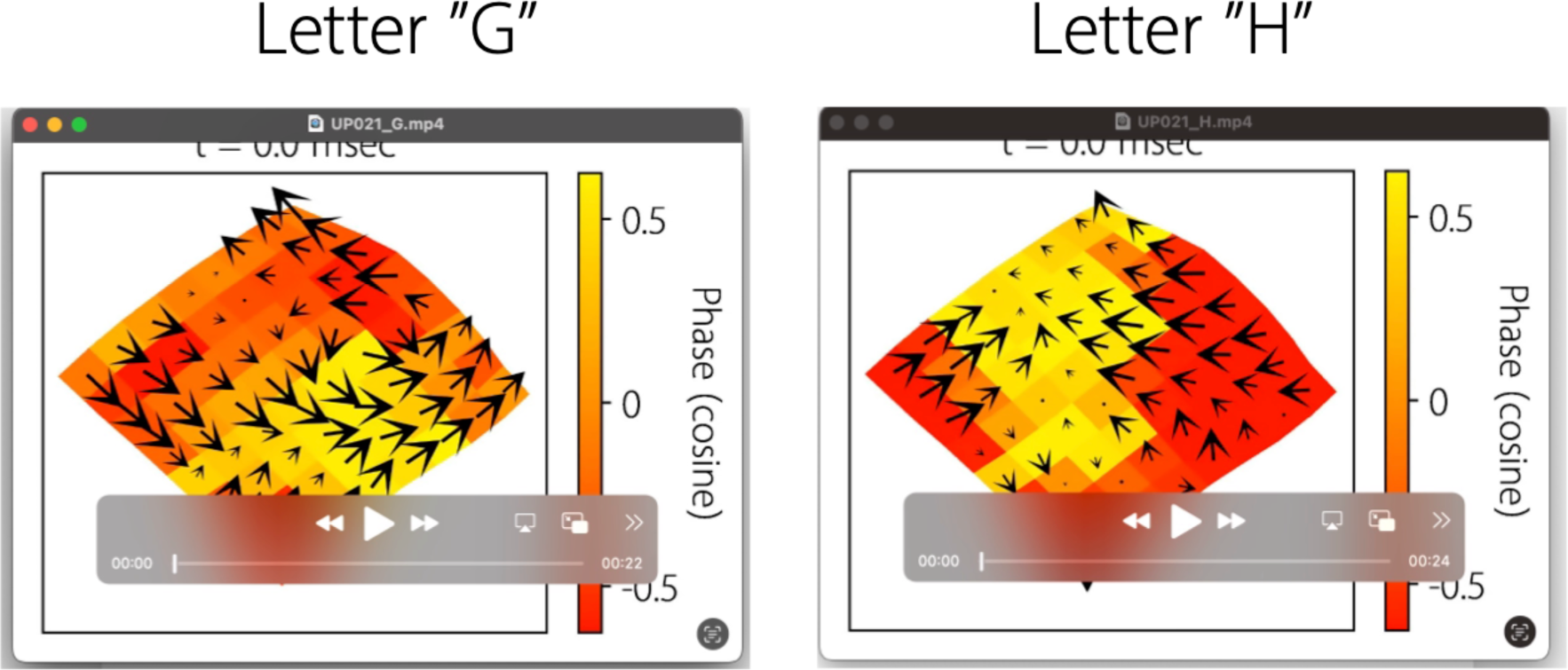
Propagation of traveling waves for two example epochs, one for the Letter “G”, and another for the Letter “H”, from patient #21, CF ∼ 12.4 Hz. Note that for the letter “G”, we observe a rotational wave, whereas for the letter “H”, we observe a sink. Shown above are snapshots of the videos. Video link: https://github.com/anupdas777/complex_traveling_waves/blob/main/UP021_G.mp4 https://github.com/anupdas777/complex_traveling_waves/blob/main/UP021_H.mp4

**Supplementary Video 3:**
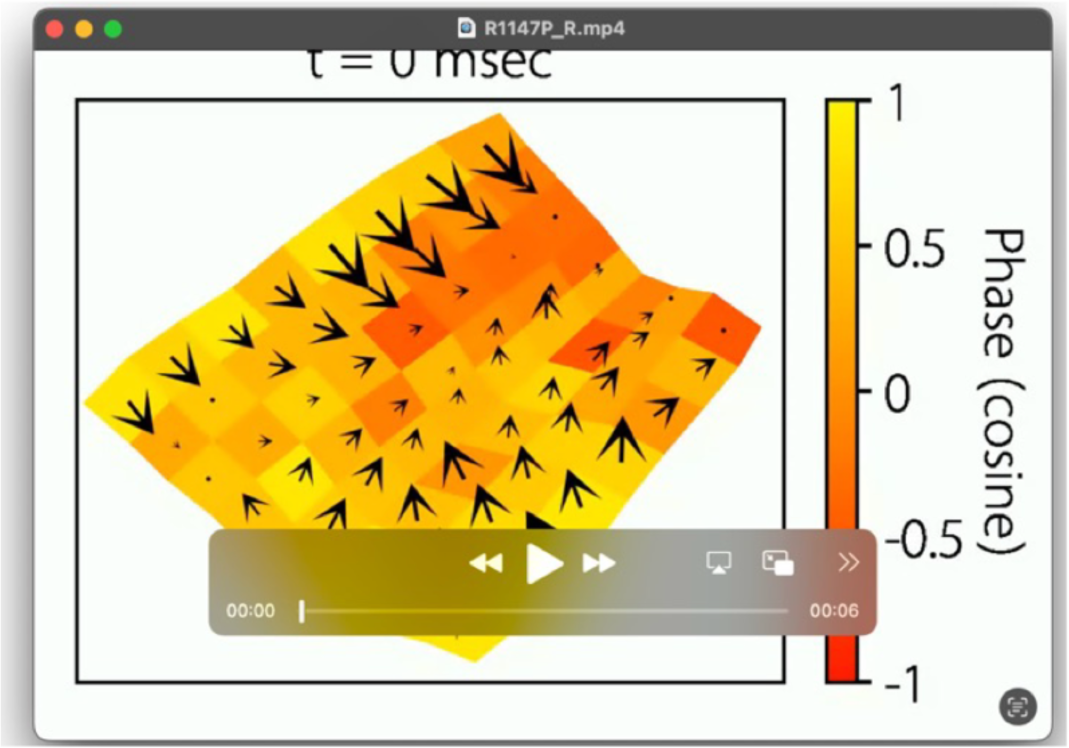
Propagation of traveling waves for an example epoch from patient #2, CF ∼ 20.6 Hz, demonstrating that rotational waves were stable at the individual epoch level (related to Figures 7A-C) Shown above is a snapshot of the video. Video link: https://github.com/anupdas777/complex_traveling_waves/blob/main/R1147P_R.mp4

**Supplementary Video 4:**
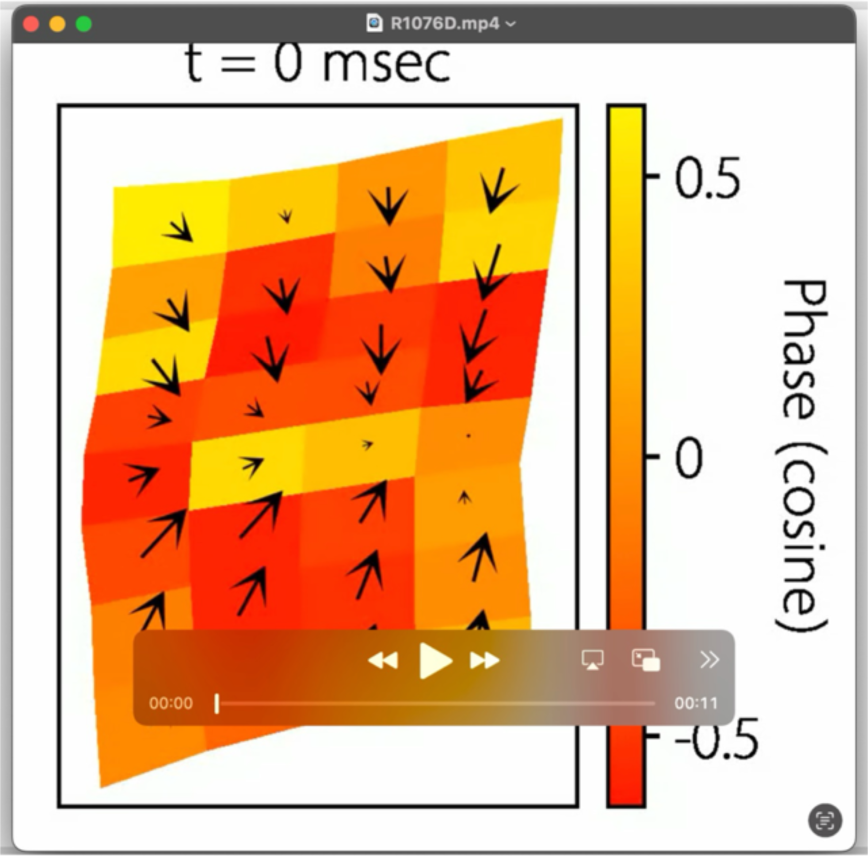
Propagation of traveling waves for an example epoch from patient #1, CF ∼ 5.6 Hz, demonstrating that concentric (expanding/contracting) waves were stable at the individual epoch level (related to Figures 8A-C) Shown above is a snapshot of the video. Video link: https://github.com/anupdas777/complex_traveling_waves/blob/main/R1076D_E.mp4

**Supplementary Video 5:**
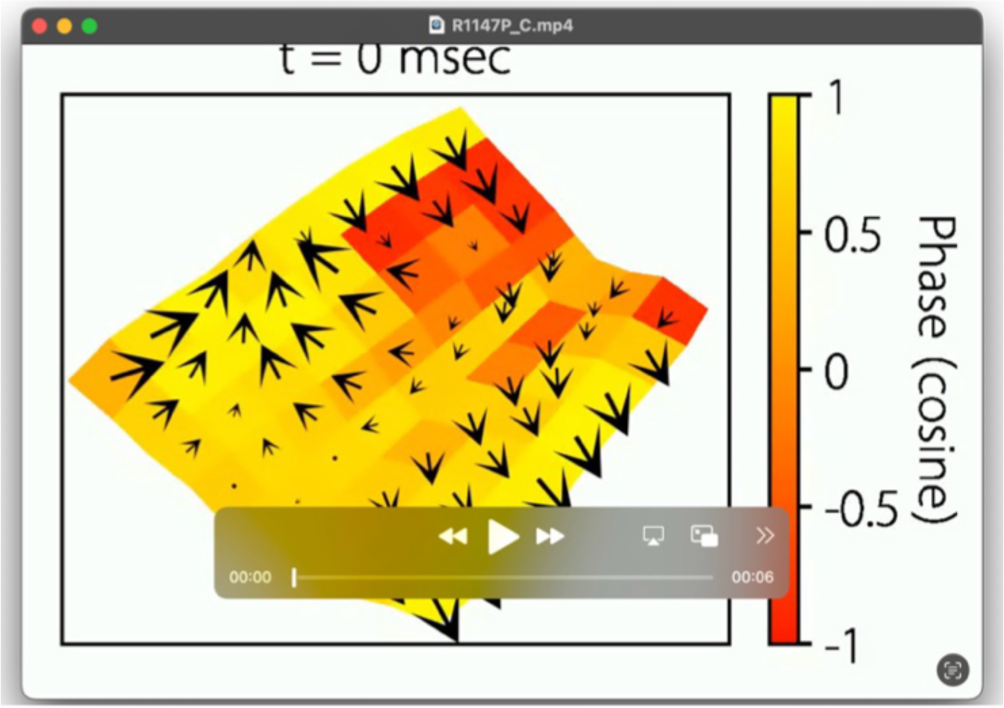
Propagation of traveling waves for an example epoch from patient #2, CF ∼ 20.6 Hz, demonstrating that complex waves were stable at the individual epoch level (related to Figures 9A-C) Shown above is a snapshot of the video. Video link: https://github.com/anupdas777/complex_traveling_waves/blob/main/R1147P_C.mp4

### Supplementary figures

**Supplementary Figure 1:**
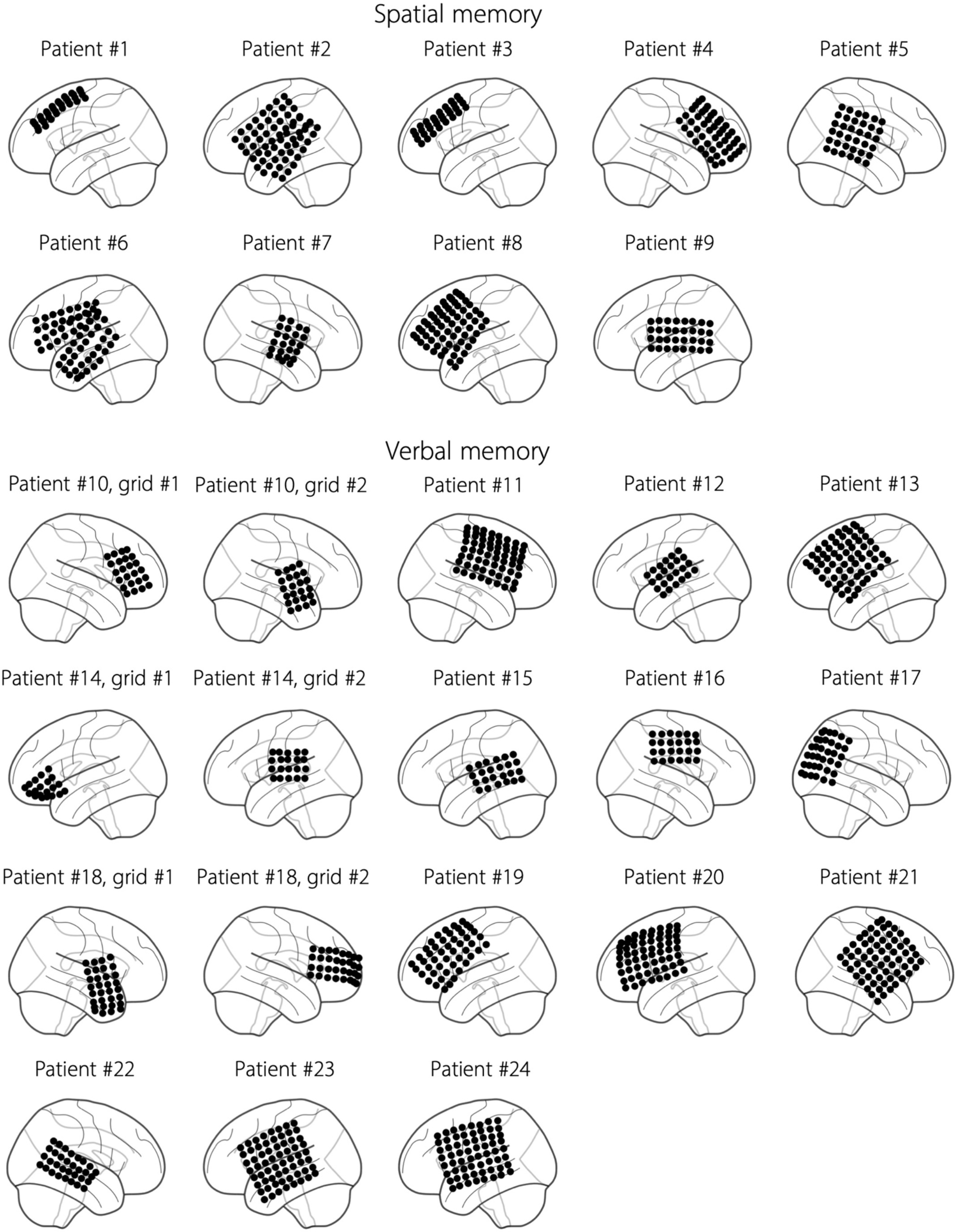
Electrode locations in the spatial (patients #1-9) and verbal (patients #10-24) memory tasks.

**Supplementary Figure 2:**
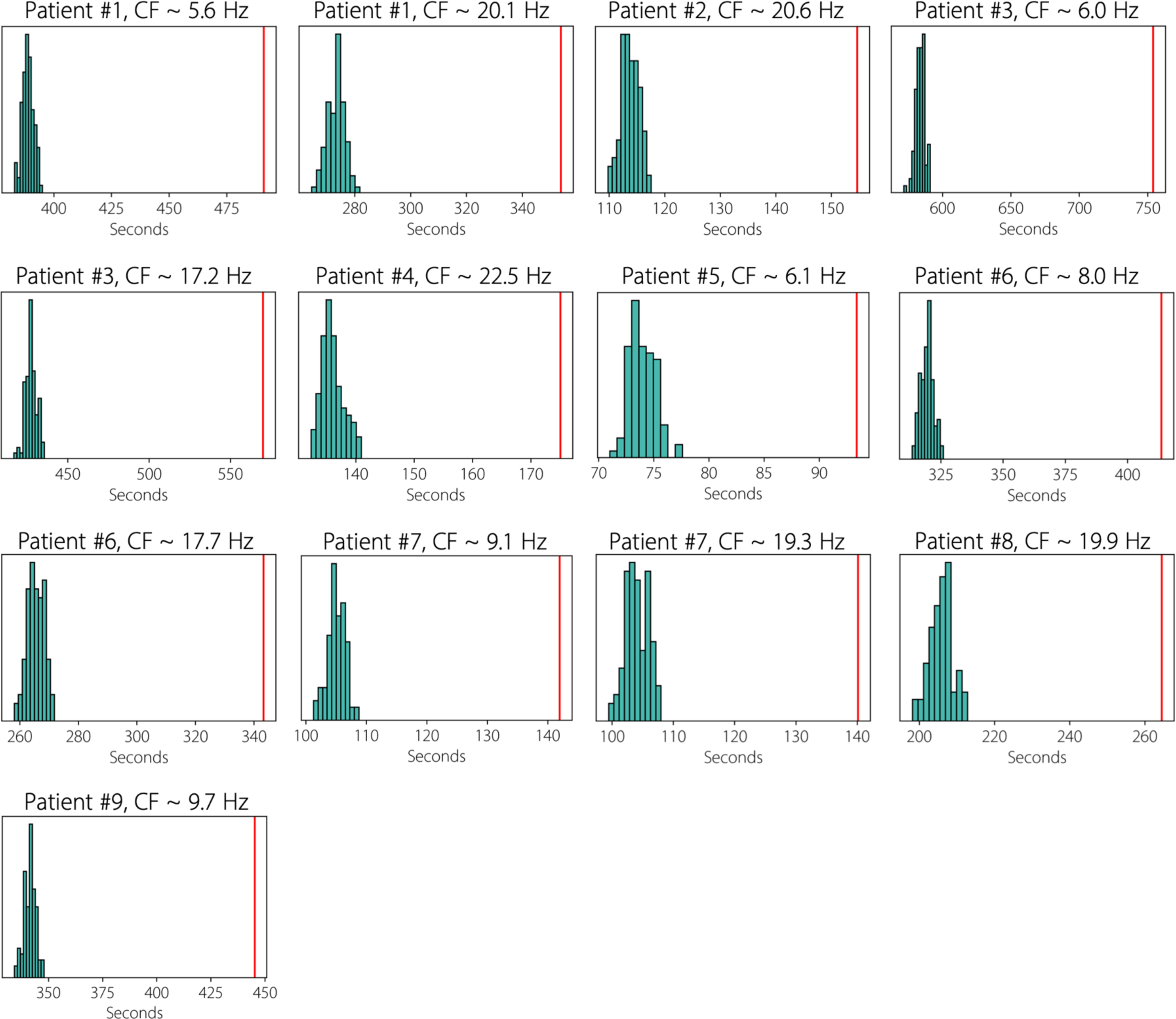
Stable epochs in the empirical data lasted significantly longer than the shuffled data, in the spatial memory task (Methods). Each panel corresponds to one cluster, with cluster frequency (CF) noted in the title of each panel, where data for the shuffled distribution is plotted in green and empirical data is plotted in red. We obtained very similar results for the verbal memory task (data not shown)

**Supplementary Figure 3:**
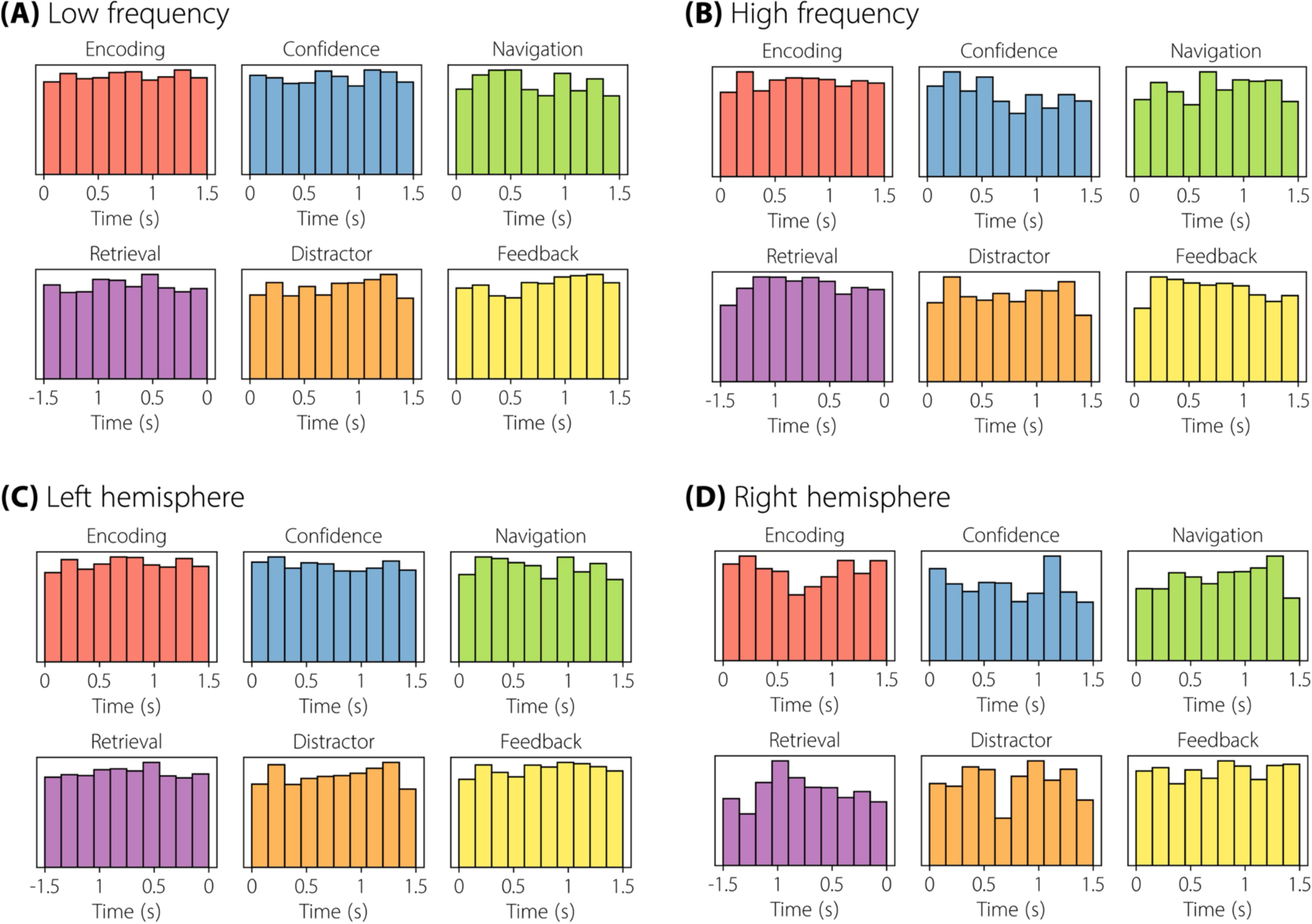
Histogram of occurrence of stable epochs across all (A) low frequency (< 12 Hz), (B) high frequency (> 12 Hz), (C) left hemisphere, and (D) right hemisphere oscillation clusters in different task periods in the spatial memory task.

**Supplementary Figure 4:**
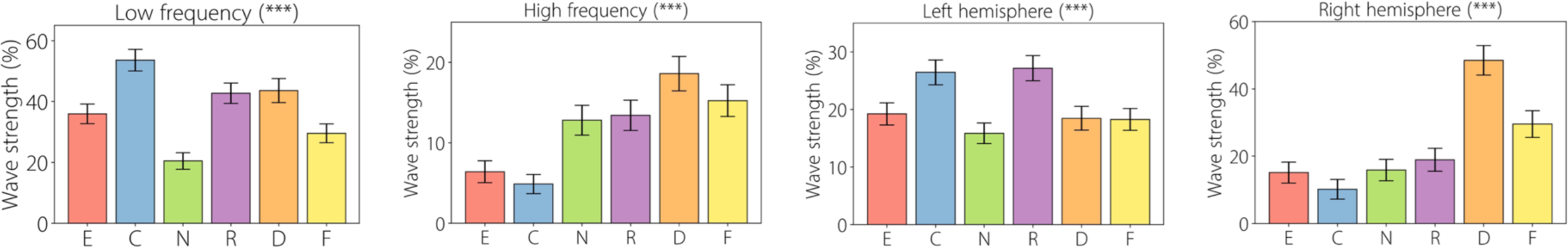
Wave strength (%) for each task period for the planar wave category in the spatial memory task, shown separately for low and high frequencies and left and right hemispheres. E: Encoding, C: Confidence, N: Navigation, R: Retrieval, D: Distractor, F: Feedback. *** *p* < 0.001 (FDR-corrected).

**Supplementary Figure 5:**
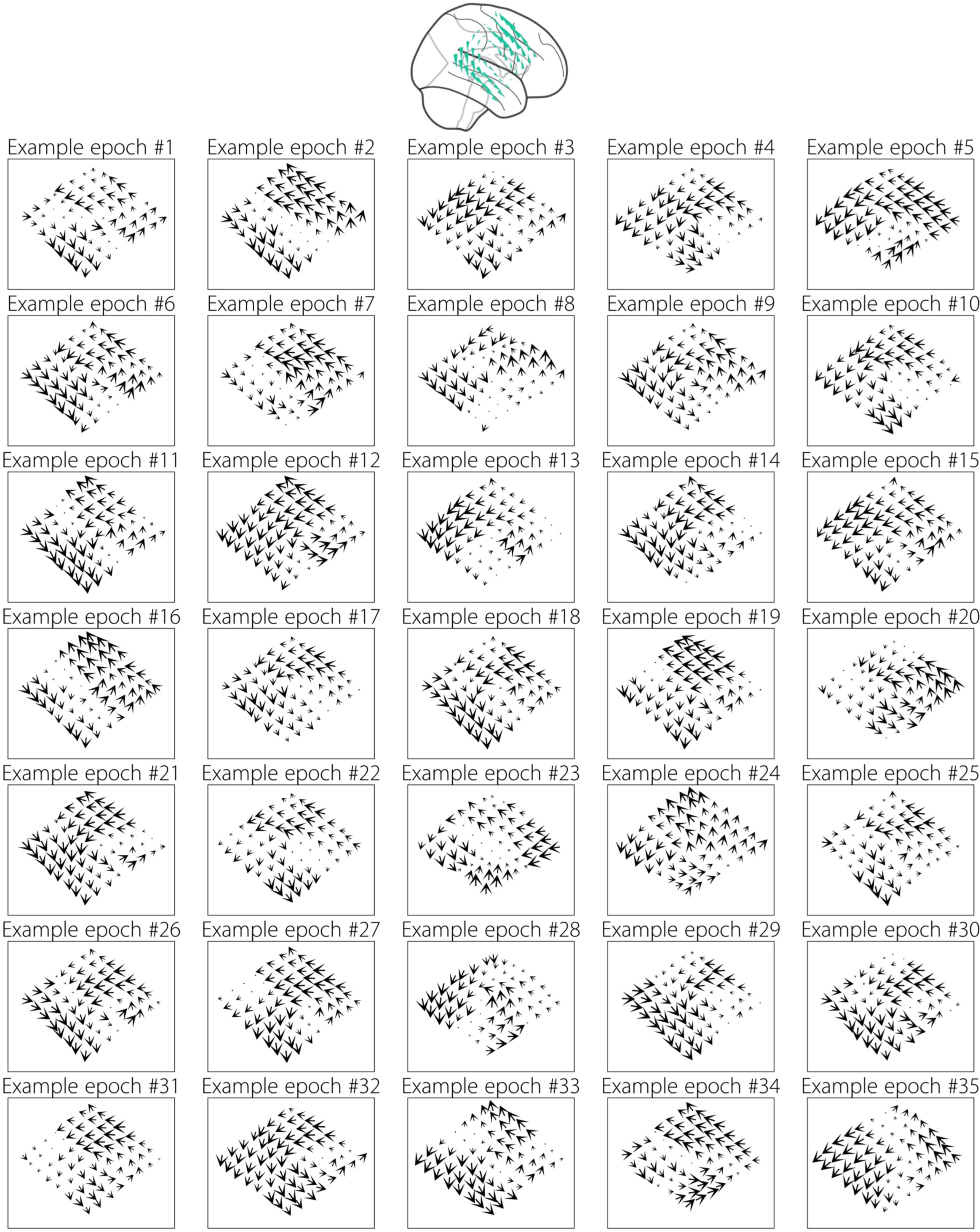
Individual epoch-level examples of rotational traveling waves for mode 2 in patient #21, CF ∼ 12.4 Hz, demonstrating that the ICA modes are present at the single-trial level.

**Supplementary Figure 6:**
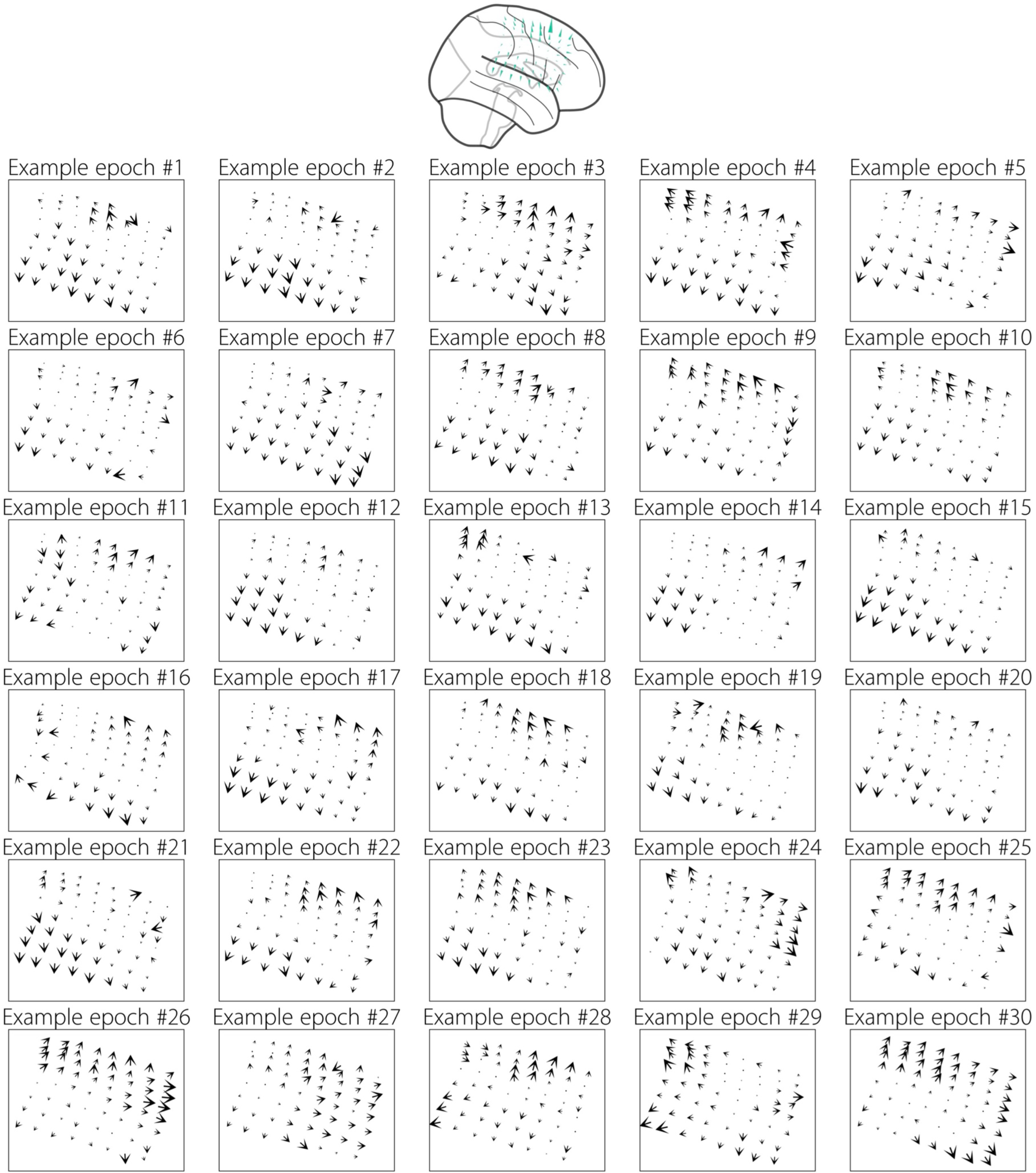
Individual epoch-level examples of expanding traveling waves for mode 2 in patient #11, CF ∼ 19.8 Hz, demonstrating that the ICA modes are present at the single-trial level.

**Supplementary Figure 7:**
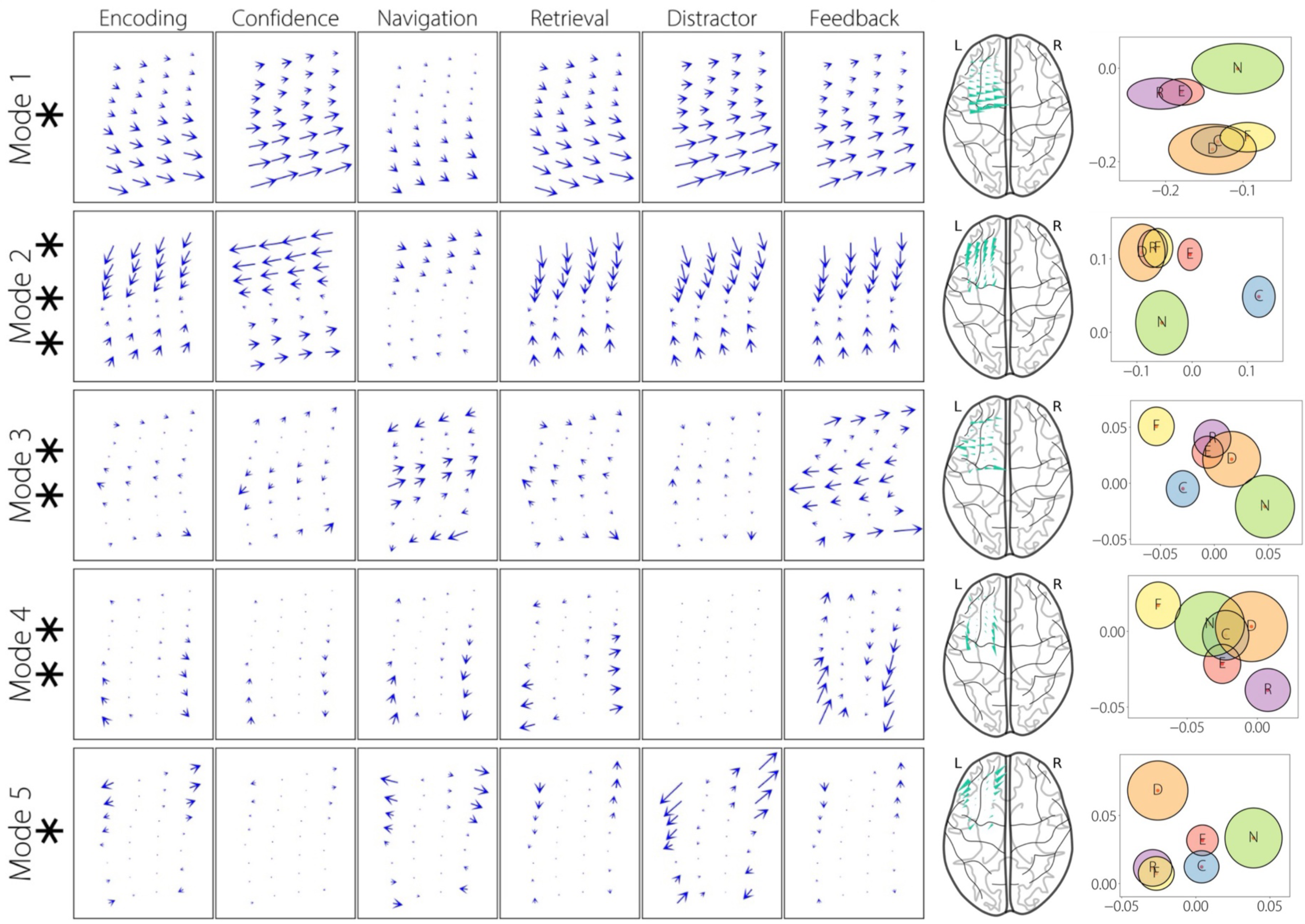
Columns 1-6: Comparing traveling waves for patient #1, CF ∼ 5.6 Hz, for the task periods encoding, confidence, navigation, retrieval, distractor, and feedback, for each mode. For each mode, shown above is the raw mode multiplied by the mean of the activation functions (or, weights) for each task period (**Methods**). Statistical significance was assessed using MANOVA (Methods). Significant modes are plotted in blue and non-significant modes are plotted in black. **Column 7: Visualization of traveling waves for each mean mode on a brain surface plot. Column 8: Visualizing the activation functions in the complex-plane for the task periods for each mode**. For each ellipse (task period), the major axis (horizontal axis) denotes the standard-error-of-the-mean (SEM) for the real-part and the minor axis (vertical axis) denotes the SEM for the imaginary part, of the activation functions. E: Encoding, C: Confidence, N: Navigation, R: Retrieval, D: Distractor, F: Feedback. *** *p* < 0.001, ** *p* < 0.01, * *p* < 0.05 (FDR-corrected).

**Supplementary Figure 8:**
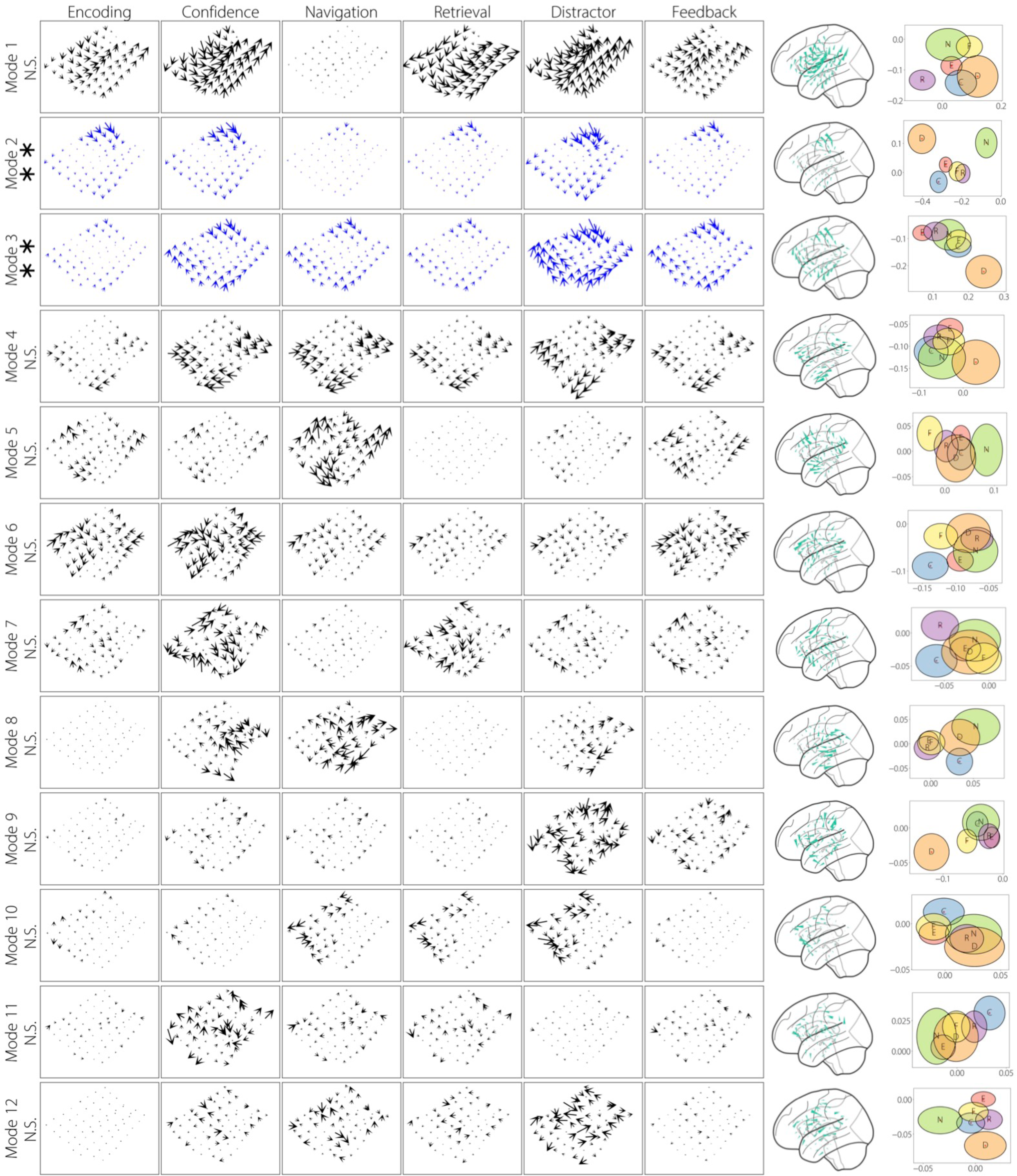
Caption similar to Supplementary Figure 7, for patient #2, CF ∼ 20.6 Hz.

**Supplementary Figure 9:**
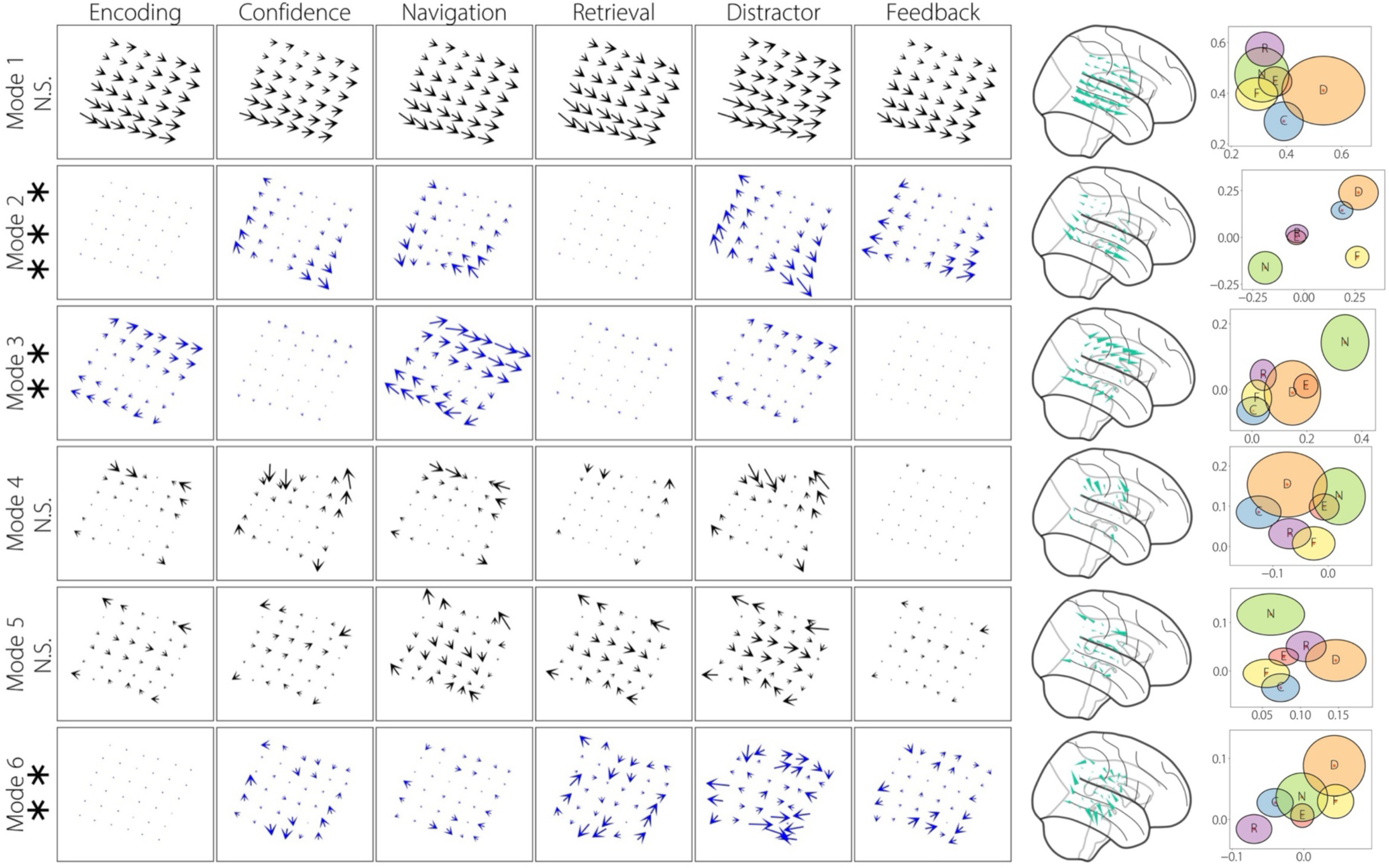
Caption similar to Supplementary Figure 7, for patient #5, CF ∼ 6.1 Hz.

**Supplementary Figure 10:**
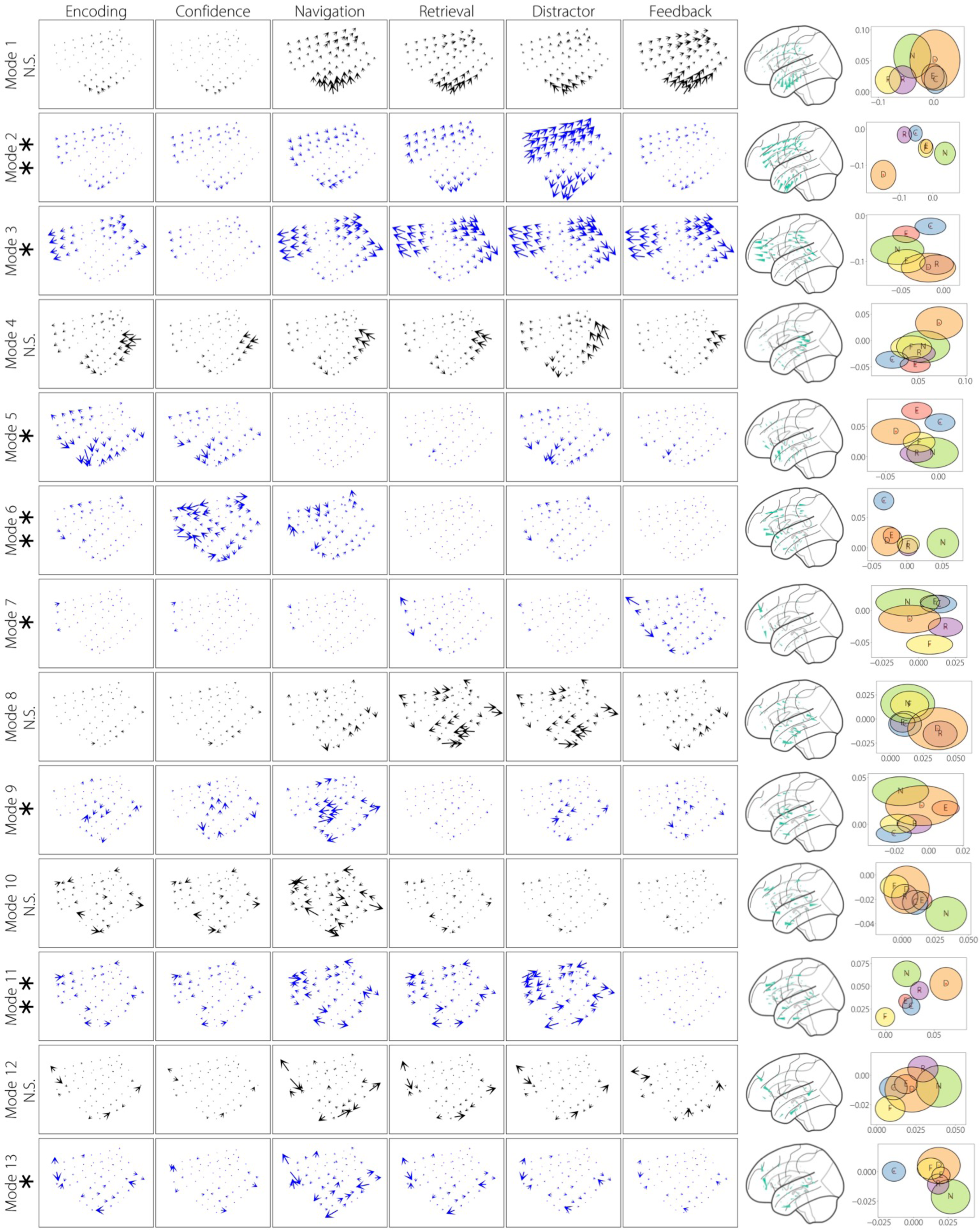
Caption similar to Supplementary Figure 7, for patient #6, CF ∼ 17.7 Hz.

**Supplementary Figure 11:**
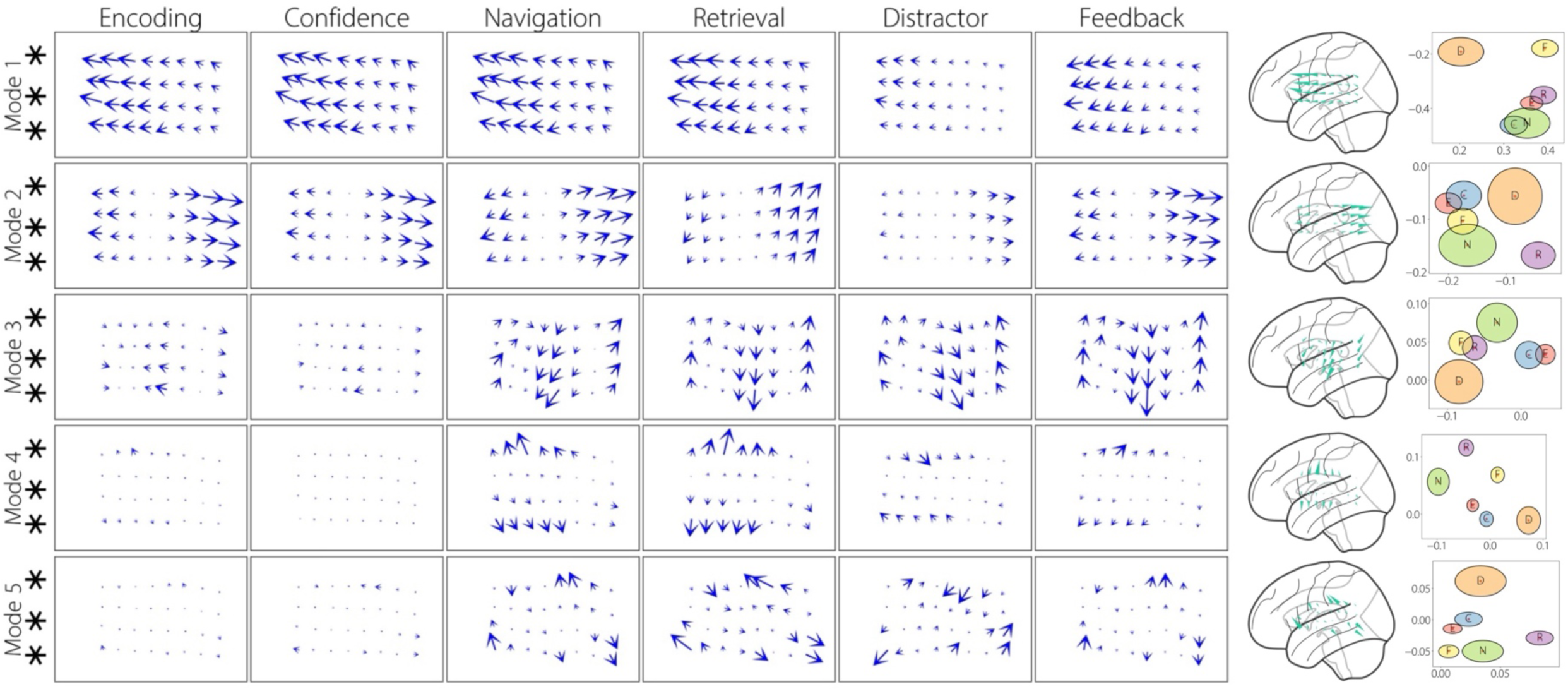
Caption similar to Supplementary Figure 7, for patient #9, CF ∼ 9.7 Hz.

**Supplementary Figure 12:**
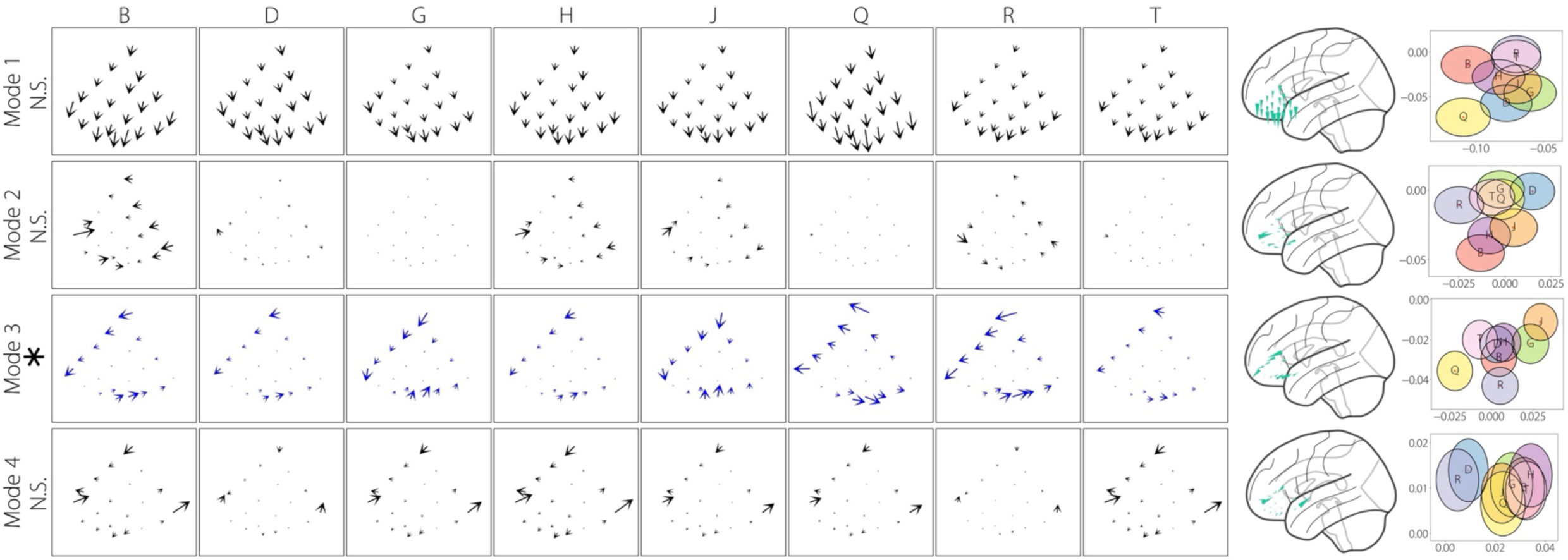
Columns 1-8: Comparing traveling waves for patient #14, grid #1, CF ∼ 25.8 Hz, for the English letters, for each mode. For each mode, shown above is the raw mode multiplied by the mean of the activation functions (or, weights) for each letter (**Methods**). Significant modes are plotted in blue and non-significant modes are plotted in black. **Column 9: Visualization of traveling waves for each mean mode on a brain surface plot. Column 10: Visualizing the activation functions in the complex-plane for the letters for each mode**. *** p < 0.001, ** p < 0.01, * p < 0.05 (FDR-corrected).

**Supplementary Figure 13:**
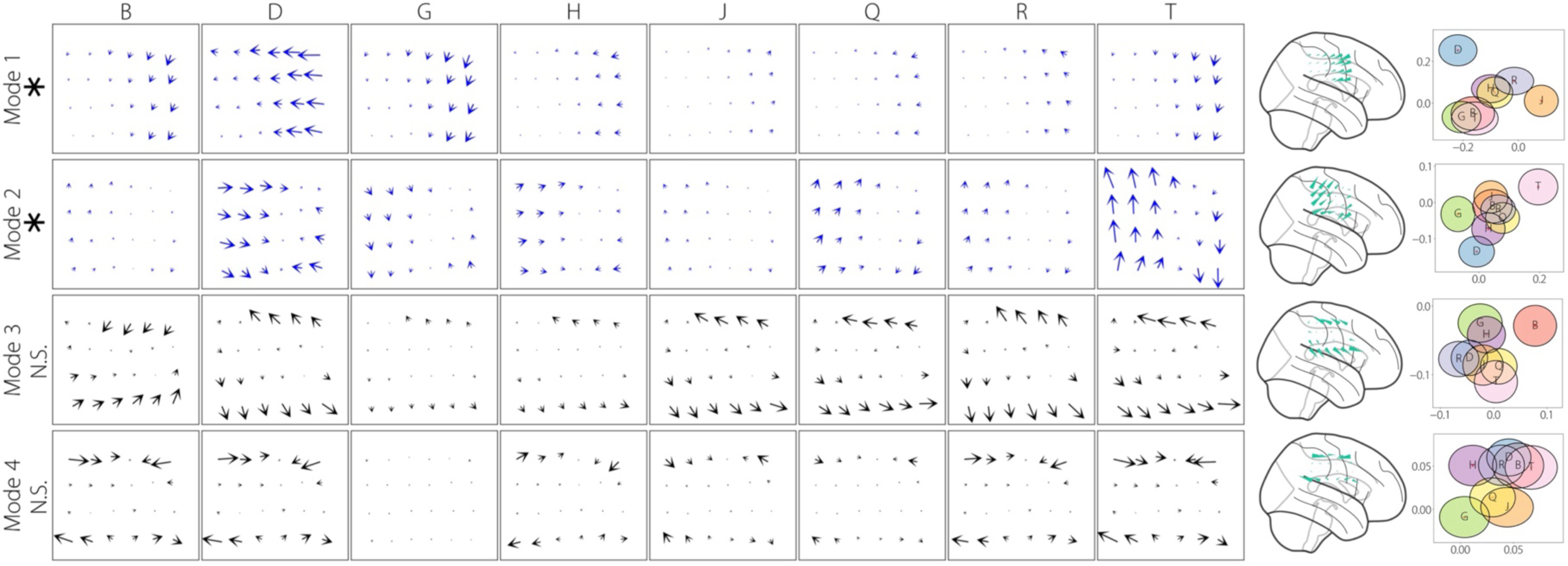
Caption similar to Supplementary Figure 12, for patient #16, CF ∼ 9.2 Hz.

**Supplementary Figure 14:**
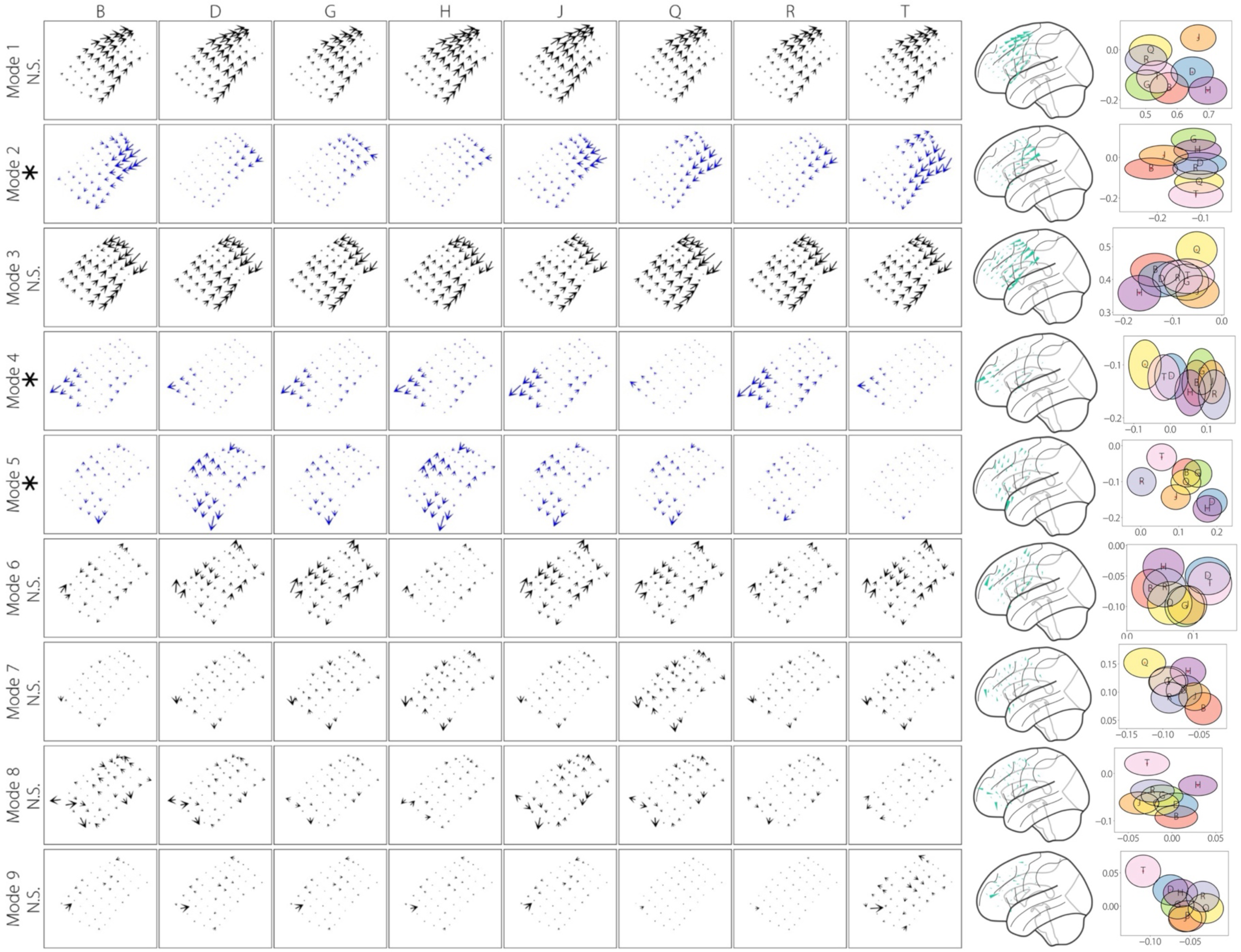
Caption similar to Supplementary Figure 12, for patient #19, CF ∼ 6.5 Hz.

**Supplementary Figure 15:**
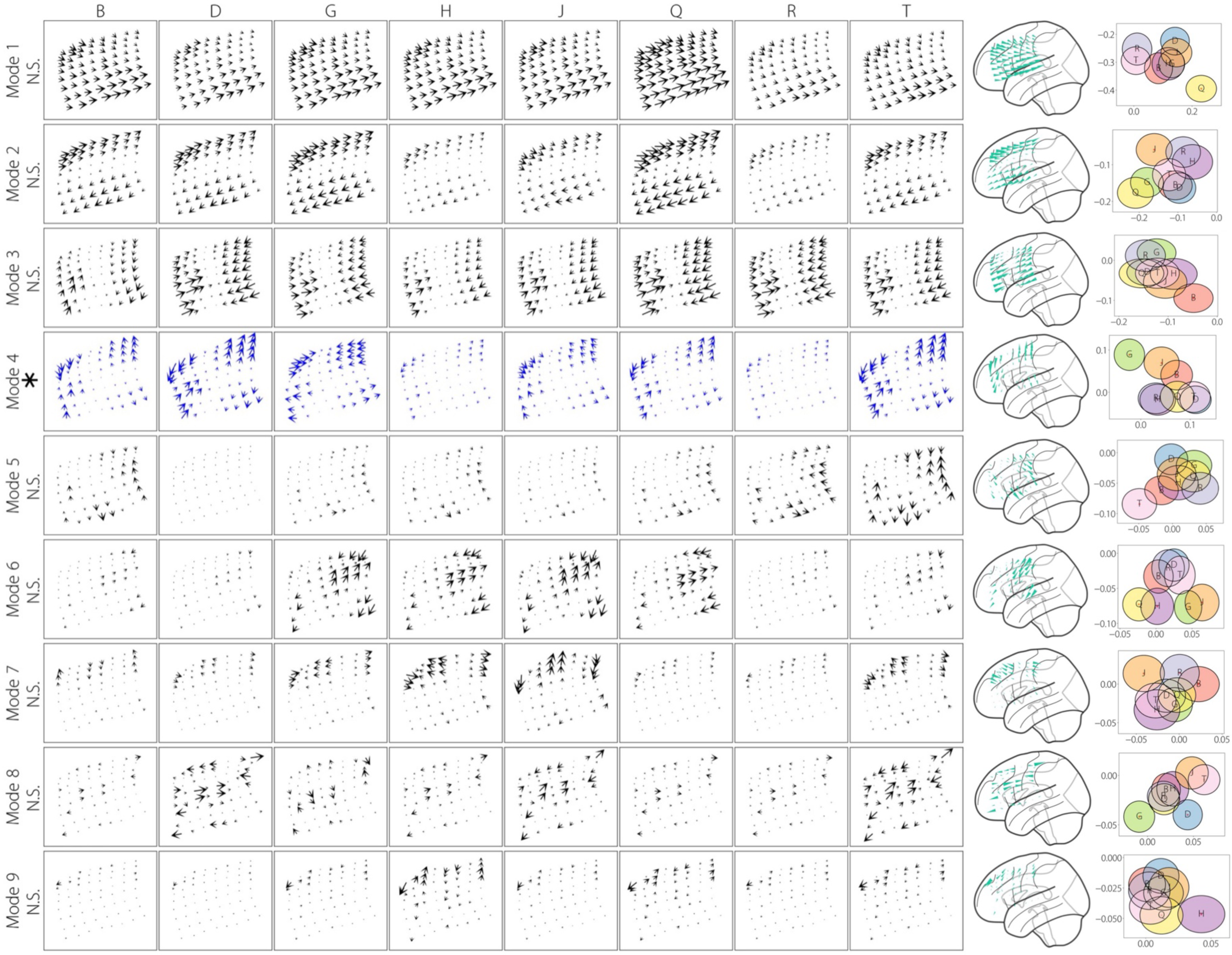
Caption similar to Supplementary Figure 12, for patient #20, CF ∼ 5.7 Hz.

**Supplementary Figure 16:**
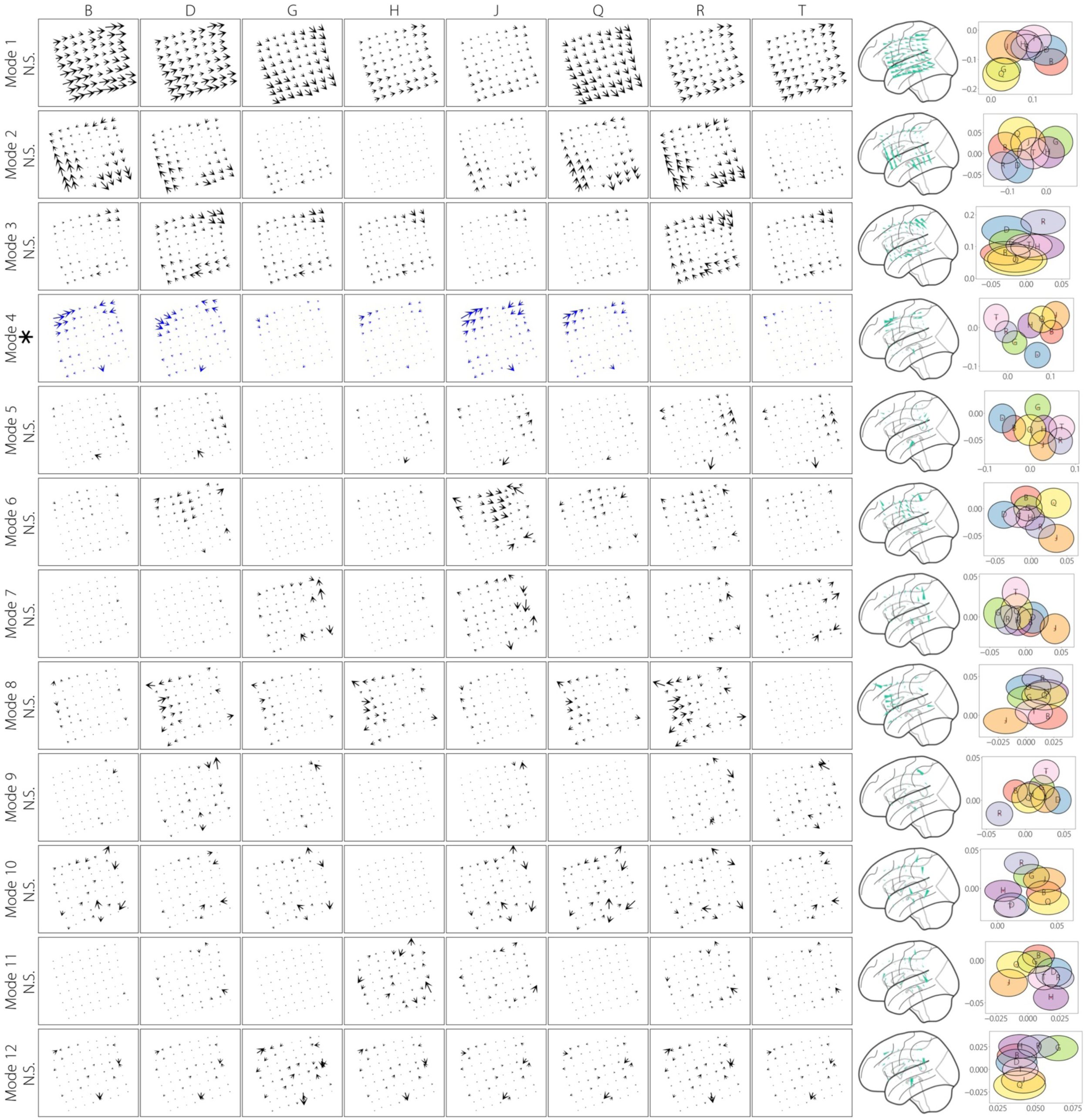
Caption similar to Supplementary Figure 12, for patient #24, CF ∼ 22.7 Hz.

